# Relating neighborhood deprivation to childhood obesity in the ABCD Study®: evidence for theories of neuroinflammation and neuronal stress

**DOI:** 10.1101/2022.06.04.494818

**Authors:** Shana Adise, Andrew T. Marshall, Eric Kan, Marybel Gonzalez, Elizabeth R. Sowell

**Author notes:** **Corresponding Author:** Elizabeth Sowell, PhD, Professor of Pediatrics | Keck School of Medicine, University of Southern California Division of Research on Children, Youth and Families | Department of Pediatrics Children’s Hospital Los Angeles 4650 Sunset Blvd., Mailstop #130 | Los Angeles, CA 90027 | |.

## Abstract

**Objective:** We evaluated whether the relationships between area deprivation (ADI), body mass index (BMI) and brain structure (e.g., cortical thickness, subcortical volume) during pre-adolescence supported the neuroinflammation (NI) and/or neuronal stress (NS) theories of overeating. The NI theory proposes that ADI causes structural alteration in the brain due to the neuroinflammatory effects of overeating unhealthy foods. The NS theory proposes that ADI-related stress negatively impacts brain structure, which causes stress-related overeating and subsequent obesity.

**Methods:** Data were gathered from the Adolescent Brain Cognitive Development^SM^ Study^®^ (9-12-years-old; n=2872, 51.3% female). Linear mixed-effects models identified brain regions that were associated with both ADI and BMI; longitudinal mediation models assessed potential causal pathways. The NI model included ADI and BMI at 9/10-years-old and brain data at 11/12-years-old. The NS model included ADI and brain data at 9/10-years-old and BMI at 11/12-years-old.

**Results:** In the NI model, BMI at 9/10-years-old positively mediated the relationship between AD and cortical thinning in the cuneus, lingual, and paracentral gyrus and larger volume of the Ventral DC at 11/12-years-old. In the NS model, cortical thinning in the lateral orbitofrontal cortex, lingual gyrus and larger volume of the Ventral DC at 9/10-years-old partially mediated the relationship between ADI and BMI at 11/12-years-old.

**Conclusion:** Greater area deprivation may indicate fewer access to resources that support healthy development, like nutritious food and non-stressful environments. Our findings provide evidence in support of the neuroinflammation and stress theories of overeating, specifically, with greater ADI influencing health outcomes of obesity via brain structure alterations.

## Introduction

Environmental resources, such as good nutrition and safety, are critical for optimal development and decreased access or opportunities for resources in the environment can negatively impact health outcomes (Hurley et al., 2016; Rosales et al., 2009). Therefore, it is not surprising that neighborhood (i.e., area) deprivation has been related to negative health outcomes, like increased risk for childhood obesity (Papas et al., 2007; Sharifi et al., 2016), and negative effects on developmental outcomes such as decreased cognitive performance and altered brain structure and function (Brito & Noble, 2014; Cockerham et al., 2017; Noble et al., 2015; Smith & Pollak, 2020; Twaits & Alwan, 2020; Vargas et al., 2020). However, despite their interconnectedness, the relationships between neighborhood deprivation, childhood obesity, and brain structural development are not well understood.

The neuroinflammation theory of overeating posits that the association between neighborhood deprivation and childhood obesity may be due to increased accessibility to unhealthy foods (Papas et al., 2007), and that repeated intake of unhealthy foods may lead to structural and functional changes in the brain due to neuroinflammation. Neuroinflammation is biochemical response to overeating unhealthy foods that is associated with brain tissue atrophy (Mullins et al., 2020). On the other hand, the neuronal stress theory of overeating suggests that the stressful impacts of living in a deprived neighborhood causes structural and functional changes in the brain, possibly due to elevated neurochemical responses (e.g., cortisol) (Barrington et al., 2014). It has been posited that overexpression of these neuroinflammation and neuronal stress markers induce structural and functional changes in the brain in regions that may facilitate and maintain overeating (Razzoli et al., 2017). Together, both of these theories suggest that neighborhood resource deprivation poses a great risk for early-onset health comorbidities (Gurnani et al., 2015) and suggest two potential mechanisms that may explain how resource deprivation facilitates and maintains overeating and subsequent obesity. However, no studies have studied the joint effects of neighborhood deprivation on obesity and brain structure. Childhood obesity continues to rise and disproportionately affects groups that have been historically economically and socially marginalized (Ogden et al., 2020), Thus, understanding how neighborhood deprivation contributes to facilitating and maintaining childhood obesity within the neuroinflammation and neuronal stress frameworks of overeating may provide optimal insight for health policy and intervention in communities that have been disproportionately affected.

Area (i.e., neighborhood) deprivation index score (ADI) is a measure of access to resources and opportunities in a given neighborhood (Kind & Buckingham, 2018). ADI incorporates aspects of the living environment such as physical infrastructure (e.g., living spaces, walkability, grocery stores, and medical resources), and social factors (e.g., education, affordability, income-to-needs, safety) that work together to influence lifestyles. Area deprivation is the result of systemic inequities in the United States and thus disproportionately impacts some populations more than others, resulting in greater health disparities in children (Singh et al., 2017). Further, area deprivation is thought to be a critical social determinant of health for child obesity. Greater deprivation is thought to increase obesogenic behaviors such as overconsumption of unhealthy food potentially due to a lack of access to healthy options or grocery stores and decreased physical activity due to a lack of access to safe, green or walkable space (Papas et al., 2007). In response to unhealthy obesogenic behaviors, neuroinflammation may occur (Mullins et al., 2020), which has been associated with downstream effects on cognitive functioning in animals (Castanon et al., 2015) and adults (Moreno-Navarrete et al., 2017). Although it has not been established that neuroinflammation is directly related to decreased cognitive performance in children, research does show a link between obesity and altered brain structure (Adise, Allgaier, et al., 2021), function (Black et al., 2014; Bruce et al., 2010), cognition and behavior (Adise, White, et al., 2021; Laurent et al., 2020; Nederkoorn et al., 2006; Verbeken et al., 2009). Moreover, area deprivation may moderate the strength of the relationship between childhood obesity and cortical thickness in regions associated with inhibitory control and executive function (Hall et al., 2021).

Furthermore, living in a neighborhood with greater area deprivation is associated with more stressors (e.g., neighborhood safety, affordability) and altered cortisol levels (Barrington et al., 2014); chronic stress has also been associated with overeating in humans and animals (Razzoli et al., 2017). Stressful environments are thought to activate the hypothalamus to induce cravings for palatable food, where food intake is proposed to downregulate negative effects of stress in the amygdala and nucleus accumbens (Razzoli & Bartolomucci, 2016). Chronic stress is also linked to altered brain structure in regions associated with cognitive control and executive functioning, learning, and memory (Smith & Pollak, 2020; Vargas et al., 2020), which may alter signaling to the hypothalamus and influence dysregulated food intake. In youth, neuronal stress may be one reason to explain why brain structure differences appear prior *to* one-year (Adise, Allgaier, et al., 2021) and two-year weight gain (Adise et al., *Under Review 2021*). Yet, to date, no studies have evaluated whether stress-related associations of living in a deprived neighborhood relate to brain structure changes that may cause obesity later in adolescence.

In this report, we assess the validity of two competing (or complimentary) theoretical models that may contribute to obesity in peripubertal youth. In the first model, we tested the neuroinflammation theory of overeating which posits that neurochemical effects (i.e., neuroinflammation) of having a higher body mass index (BMI) at 9/10-years-old, possibly due to overeating of unhealthy foods, mediates the association between area deprivation at 9/10-years-old, a proxy for access to health promoting behaviors (e.g., good food, safe exercise spaces), and brain structure variation at 11/12-years-old (**Fig1A**). Although research has shown that a lack of access to good food is related to greater BMI (Papas et al., 2007), and that greater BMI is related to brain structure variation (Adise et al., 2021), no studies have assessed these relationships in tandem. In the second model, we tested the neuronal stress theory of overeating, which hypothesizes that structural variation at 9/10-years-old, possibly due to the stress-related effects of living in an area deprived of resources, mediates the relationship between area deprivation at 9/10-years-old and BMI at 11/12-years-old by triggering stress-induced overeating (**Fig1B**). Data were curated from the Adolescent Brain Cognitive Development Study (ABCD Study^®^). Although the ABCD Study^®^ collects various MRI modalities, we focused on cortical thickness and subcortical volume, as prefrontal cortical thinning has cross-sectionally showed a relationship with ADI and BMI at 9/10-years-old (Hall et al., 2021), and much of food intake is regulated by subcortical regions (Berthoud, 2012). As some indices of area deprivation can be modified to reduce social inequities leading to health disparities, understanding how it relates to obesity development and maintenance may be crucial for health policy and intervention.

**Fig1.**
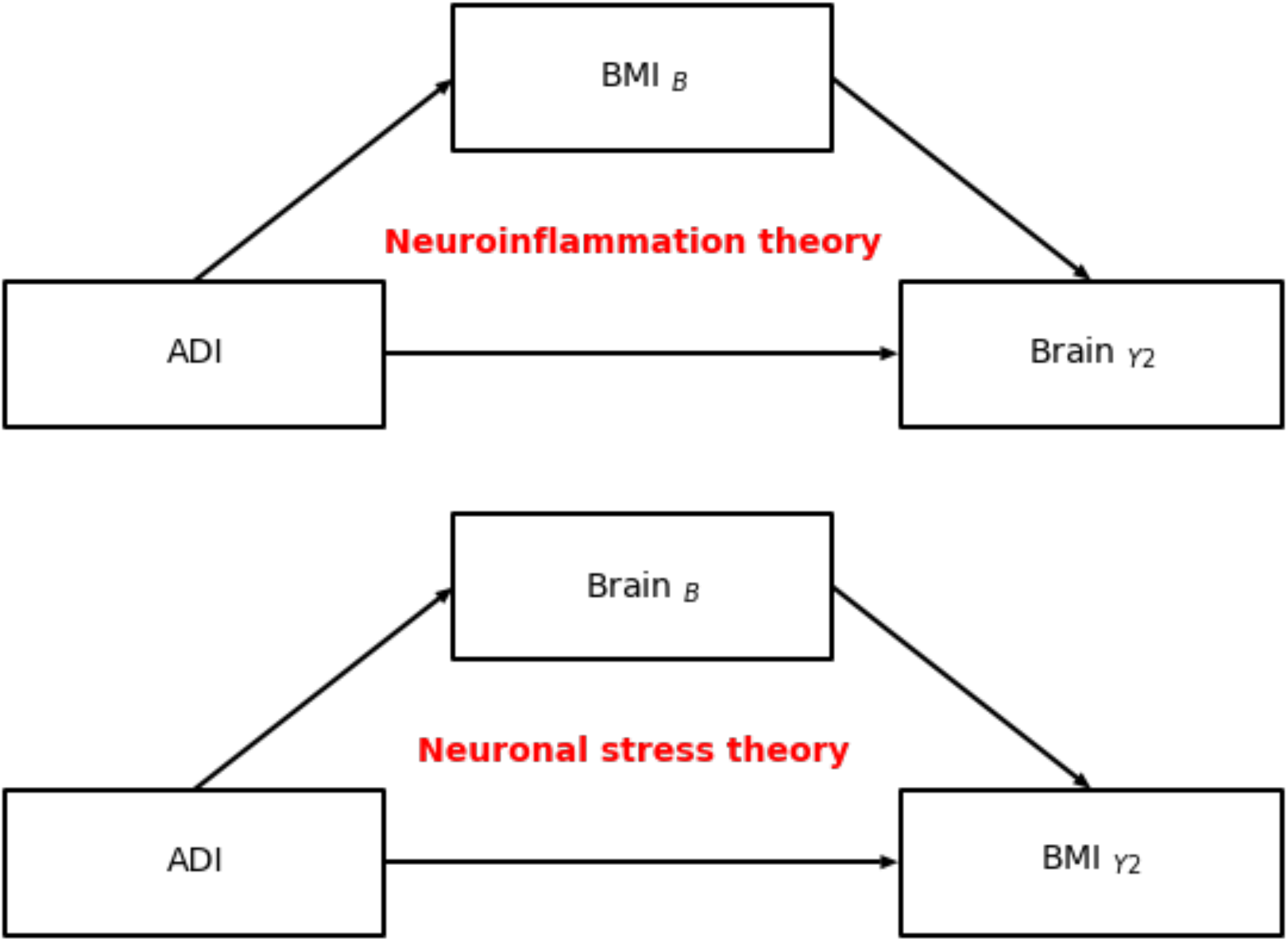
A schematic overview of the neuroinflammation and neuronal stress theories of overeating. B = baseline (aged 9/10-years-old). Y2 = two-year follow up (aged 11/12-years-old).

## Methods

### Study design

The ABCD Study^®^ is a 21-site 10-year longitudinal cohort study (Auchter et al., 2018; Uban et al., 2018; Volkow et al., 2018). The current manuscript presents data from the 3.0 release (*n*_baseline_=11,878; *n*_year1_=11,235, *n*_year2_ = 6,571 youth) with neuroimaging and anthropometric data (acquired at baseline and the two-year follow-up). All data were collected prior to the onset of the coronavirus pandemic. Caregivers and youth provided written consent. A centralized institutional review was approved by the University of San Diego.

### Exclusion criteria

A full list of exclusion criteria determined by the ABCD Study^®^ are listed elsewhere (Garavan et al., 2018), but additional exclusion criteria were applied in order to obtain a sample that was acceptable for these analyses and hypotheses. Youth were excluded if they met the following at any of the time points (e.g., baseline, one-year or two-year follow-up): (1) Underweight (according to the Center for Disease Control’s (CDC’s) age-sex-height-weight-specific growth curves (Kuczmarski et al., 2002)) possibly due to restrictive eating or medical issues; (2) taking medications known to alter food intake (e.g., antipsychotics, insulin); (3) had caregiver report of neurological, psychiatric, or learning disabilities; (4) met diagnostic criteria for eating disorders (e.g., anorexia, binge eating disorder) as assessed by the caregiver-reported Kiddie Schedule for Affective Disorders and Schizophrenia (Kaufman et al., 1997); (5) mislabeled sex-assigned at birth and/or mismatched sex-specific pubertal questionnaires or transgender youth (i.e., due to sex-specific effects on brain function); (6) height measurement error (e.g., height year 2 < baseline); (7) invalid residential address (necessary for ADI metrics); (8) failed FreeSurfer segmentation; (9) failed T_1_ quality control; and/or (10) missing ROI or covariate tabulated data from the National Institutes of Mental Health databases. Siblings were excluded to avoid issues with independence. The final sample consisted of 2,872 youth (**Table1**).

**Table 1.** Participant characteristics. Baseline was assessed when the youth were 9-to 10-years-old. Y2= two-year follow up (age 9/10-years-old). BMI=Body mass index. ADI=area deprivation index. AIAN/NHPI=American Indian, Alaskan Native/Native Hawaiian and Pacific Islanders. HS=high school; GED=Generalized education diploma. BA=Bachelor’s degree. SD=standard deviation.

| Variable | Mean | SD |
| --- | --- | --- |
| Age |  |  |
| *Baseline* | 119.1 | 7.2 |
| *Y2* | 142.9 | 7.5 |
| Puberty |  |  |
| *Baseline* | 2 | 0.8 |
| *Y2* | 2.7 | 1.0 |
| BMI |  |  |
| *Baseline* | 19.1 | 3.7 |
| *Y2* | 21 | 4.5 |
| ADI score | 37.3 | 26.2 |
|  | **n** | **%** |
| Sex |  |  |
| *Male* | 1473 | 51.3 |
| *Female* | 1399 | 48.7 |
| Race |  |  |
| *White* | 1998 | 69.6 |
| *Black* | 336 | 11.7 |
| *Asian* | 69 | 2.4 |
| *AIAN/NHPI* | 28 | 1.0 |
| *Other* | 140 | 4.9 |
| *Multi-race* | 301 | 10.5 |
| Ethnicity |  |  |
| *Hispanic* | 598 | 20.8 |
| *Non-Hispanic* | 2274 | 79.2 |
| Education |  |  |
| *<HS* | 101 | 3.5 |
| *HS/GED* | 212 | 7.4 |
| *Some College* | 723 | 25.2 |
| *BA degree* | 764 | 26.6 |
| *Postgraduate degree* | 1072 | 37.3 |
| Baseline Weight Class |  |  |
| *Healthy Weight* | 1899 | 66.1 |
| *Overweight* | 489 | 17.0 |
| *Obese* | 484 | 16.8 |
| Y2 Weight Class |  |  |
| *Healthy Weight* | 1810 | 63.0 |
| *Overweight* | 522 | 18.2 |
| *Obese* | 540 | 18.8 |

### Anthropometries

Yearly height (nearest 0.1inch) and weight (nearest 0.1lb) assessments (measured twice, a third was collected in cases of large discrepancy) were gathered by a trained researcher. The closest two measurements were averaged and converted into BMI (kg/m^2^) and BMI percentiles according to the CDC’s sex-age-height-weight specific growth charts (Kuczmarski et al., 2002) for clinical interpretations, only as they are prone to several biases (Hendrickson & Pitt, 2021; Palmer et al., 2021).

### Pubertal assessment

Puberty was assessed via caregiver and self-report sex-specific questionnaires and then averaged. Scores were converted into sex-specific Tanner staging categories (1=Prepubertal, 2=Early puberty; 3=Mid puberty; 4=Late puberty; 5=Postpubertal).

### Demographic assessments

The caregiver reported the child’s race, ethnicity, date of birth, and sex at birth. Race had 22 options, which were collapsed into six groups: White, Black, Asian, American Indian, Alaskan Native/Native Hawaiian, Pacific Islander (AIAN/NHPI), Other and multi-race. Ethnicity was assessed with two options: Hispanic or Non-Hispanic. Age at each visit was recorded in months. Highest household education was assessed by caregiver report across 29 education levels and collapsed into five groups: <High school (HS), HS/Generalized Education Diploma, Some college, Four-year degree (Bachelor’s degree), Postgraduate education.

### Area Deprivation Index (ADI)

ADI is a neighborhood atlas deprivation composite score based on 17 factors (e.g., income, education, housing) from the American Community Survey. Higher values indicate higher disadvantage. ADI was calculated by the DAIRC in R (https://github.com/ABCD-STUDY/geocoding/blob/master/Gen_data_proc.R) (Fan et al., 2021).

### Neuroimaging Measures

#### Image acquisition and preprocessing

Exact details of the ABCD Study^®^ MRI data acquisition and analyses are published elsewhere (Casey et al., 2018; Hagler et al., 2019). MRI data were collected with 29 scanners, as some sites had multiple MRI acquisition centers. Youth underwent a T_1_- and T_2_-weighted MRI, diffusion tensor imaging, resting state MRI, and three functional MRI scans. The current manuscript focuses on the T_1_-weighted structural, imaging acquisition. After preprocessing, cortical data were surface projected and then parcellated with Freesurfer using the Desikan Atlas (Hagler et al., 2019), which consists of 68 regions of interest (ROIs). In addition, 16 subcortical ROIs were also parcellated from the volumetric data. ROI estimates (e.g., mean cortical thinning, total gray matter volumes) were made available through the tabulated data release. ROI estimates were averaged across hemispheres (e.g., left, right), for a total of 34 cortical and 8 subcortical ROI estimates.

### Statistics

#### Linear Mixed-Effects Modeling

Multicollinearity issues were assessed by a variance inflation factor, and winsorization was implemented to normalize outliers. Brain regions that were significantly related to both BMI and ADI were determined by implementing linear random mixed effects models with the Python package *statsmodel* (Seabold & Perktold, 2010). The models included corrections for sex, puberty (at baseline) highest household education, race, and ethnicity. Independent variables were standardized. Intracranial volume and age were not included due to multicollinearity issues. Random effects accounted for variability across scanners. Models 1-4 (below) were corrected using the Benjamini-Hochberg approach.

##### Neuroinflammation models

Model 1: ROI_Y2_ ~ BMI_B_ + puberty + sex + Race + Ethnicity + education + (1|Scanner ID)

Model 2: ROI_Y2_ ~ ADI+ puberty + sex + Race + Ethnicity + education + (1|Scanner ID)

##### Neuronal stress models

Model 3: BMI_Y2_ ~ ROI_B_ + puberty + sex + Race + Ethnicity + education + (1| Scanner ID)

Model 4: ADI ~ ROI_B_ + puberty + sex + Race + Ethnicity + education + (1|Scanner ID)

### Mediation analyses

Longitudinal mediation analyses controlled for covariates were conducted in Python with *Pyprocessmacro* (https://github.com/QuentinAndre/pyprocessmacro). Continuous variables were mean centered. Significance of the indirect effect was analyzed by conducting bootstrapping to generate bias-corrected 95% confidence intervals (CI; 10,000 samples).

## Results

### Demographics

Among the 2,872 youth included in analyses, 51.3% were female, 69.6% were White, and 79.2% were not Hispanic (**Table1**). Most youth had at least one parent with a Bachelor’s degree or higher (63.9%). The sociodemographic composition of the sample analyzed reflected the composition of the larger ABCD Study^®^ sample (Garavan et al., 2018), while the rates of overweight and obese youth were comparable to national estimates at baseline (66.1% healthy weight) and at the two-year follow-up (63% healthy weight) (Ogden et al., 2020). There was a wide range of ADIs (**Fig2A**). Partial correlations controlled for the covariates of interest showed that ADI was positively related to BMI at baseline (*r*=0.1, *p*<0.001, **Fig2B**) and the two-year follow-up (*r*=0.1, *p*<0.001, **Fig2C**).

**Fig2.**
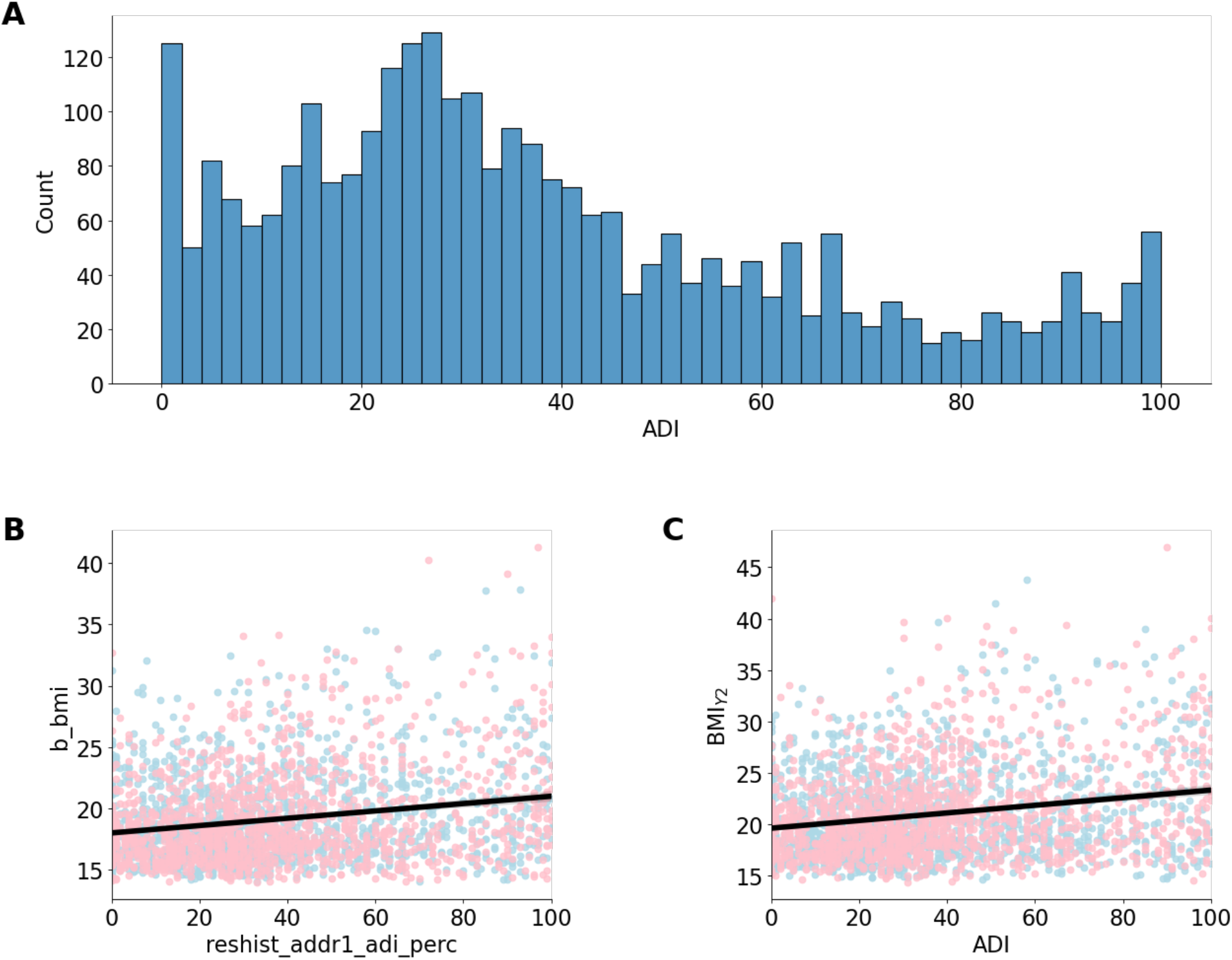
**A)** Distribution of area deprivation index (ADI; higher values = greater neighborhood deprivation). **B** and **C**) The relationship between BMI and ADI at baseline, 9/10 years-old (B) and the two-year follow up (Y2), 11/12 years-old colored by sex assigned at birth (blue=male; pink=female). The black line indicates the regression fit collapsed across sex (*r*=0.1, *p*<0.001 for both time points).

### Neuroinflammation theory of overeating

At 11/12-years-old, three cortical thickness ROIs (e.g., cuneus, lingual gyrus, paracentral gyrus) and one subcortical ROI (e.g., Ventral diencephalon [Ventral DC]) were significantly retrospectively associated with ADI and BMI at 9/10-years-old (see **Tables 2–3**).This suggest that these relationships may be driven by the environmental effects such as a lack of access to healthy foods. BMI at 9/10-years-old partially mediated the associations of ADI at 9/10-years-old on brain structure at 11/12-years-old in the cuneus, lingual gyrus, paracentral gyrus, and Ventral DC (**Fig3A**). ADI was positively associated with BMI at 9-to 10-years-old *(a* path, **Fig3B**) but negatively associated with brain structure at 11/12-years-old (C path, total direct effect, **Fig3B**). BMI at 9/10-years-old was negatively associated with cortical thinning in the cuneus, lingual, and paracentral gyri at 11/12-years-old but positively associated with subcortical volumes of the Ventral DC at 11/12-years-old *(b* path, **Fig3B**).

**Fig3.**
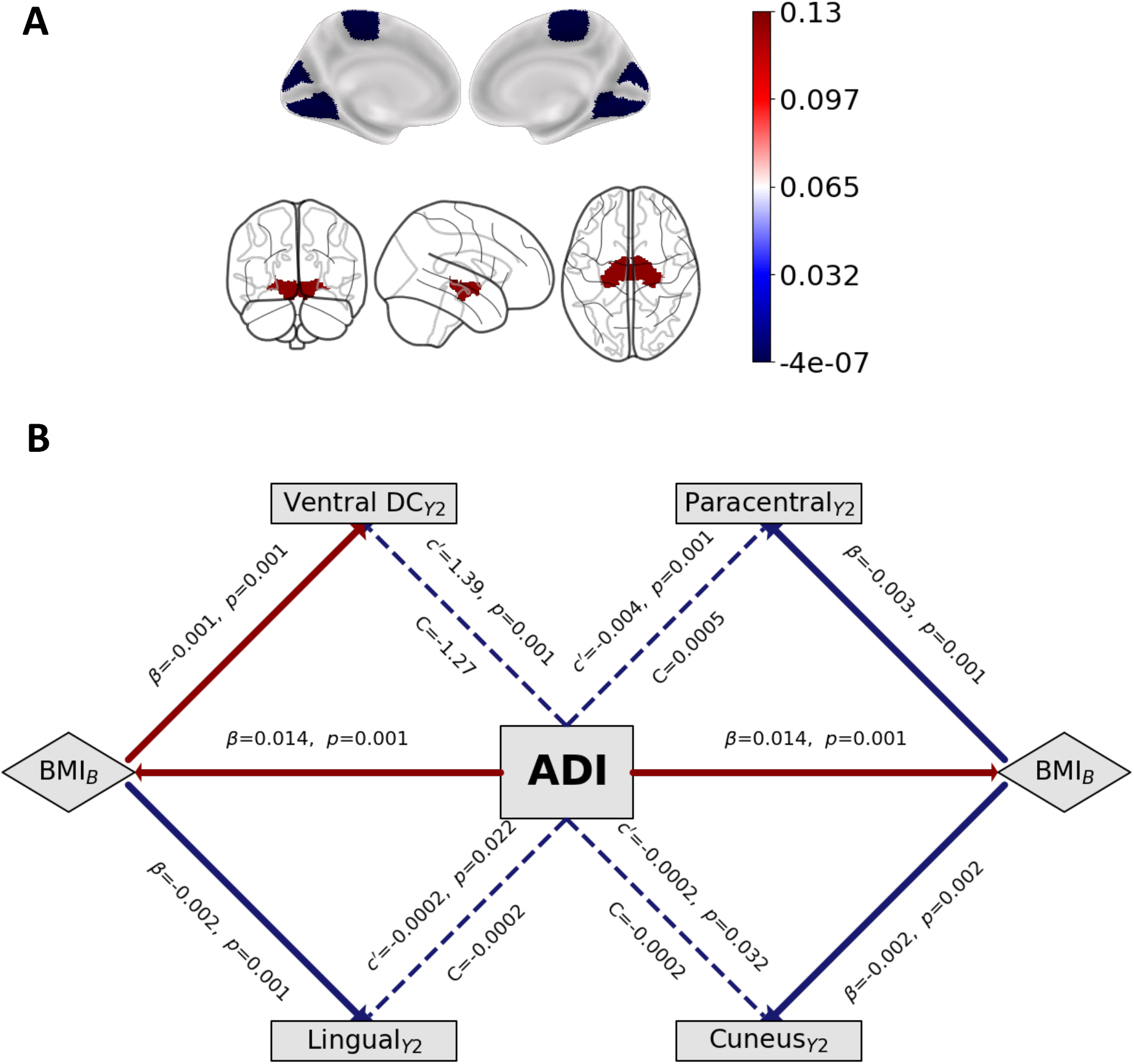
Testing the causal pathway of the neuroinflammation theory of overeating. Mediating effects of BMI at 9/10-years-old on ADI and brain structure at 11/12-years-old. **A)** Visual representation of the brain regions that showed mediating effects colored by the strength of the indirect association. **B)** Mediation models where the colored boxes reflect the strength (and direction) of the indirect effects. B=baseline (aged 9/10-years-old). Y2=two-year follow-up (aged 11/12-years-old). ADI = area deprivation index. BMI = body mass index. Total effects are represented by *c’*, direct effects of ADI are represented by C while *a* and *b* values refer to the association of ADI on BMI and BMI on brain structure, respectively. All *a, b, c*, and *c’* values are unstandardized regression coefficients. Significance testing was carried out by bias-corrected bootstrapping (n=10,000) 95% confidence intervals.

**Table 2.** Parameter estimates from the neuroinflammation model assessing the associations between ADI and BMI at 9/10-years-old and cortical thickness at 11/12-years-old. Linear random mixed effects covaried for puberty, sex, Race/ethnicity and education with a random effect of scanner type to identify ROIs at 11/12-years-old that were significantly associated with ADI and BMI at 9/10-years-old. Correction was conducted separated by model (e.g., ADI, BMI). ^a^survived multiple correction testing using the Benjamini-Hochberg method. Salmon-colored rows highlight the ROIs that showed significant associations with BMI and ADI. Blue rows highlight regions that were significant but there was no overlap between BMI and ADI. Gray rows had no significant associations.

| ROI (11-12-yrs) | Neuroinflammation model |  |  |  |  |  |
| --- | --- | --- | --- | --- | --- | --- |
|  | Associations with BMI at 9-10-yrs |  |  | Associations with ADI at 9-10-yrs |  |  |
|  | Beta | 95%CI | *p* | Beta | 95%CI | *p* |
| Anterior cingulate (Caudal) | 0.01 | [-0.03,0.05] | 0.721 | 0 | [-0.04,0.05] | 0.865 |
| Anterior cingulate (Rostral) | -0.01 | [-0.05,0.03] | 0.558 | 0.04 | [-0.01,0.08] | 0.119 |
| Caudal middle frontal | -0.03 | [-0.06,0.01] | 0.134 | -0.02 | [-0.06,0.03] | 0.457 |
| Cuneus | -0.05 | [-0.09,-0.01] | 0.01*a | -0.11 | [-0.15,-0.06] | <0.001***a |
| Entorhinal | -0.04 | [-0.08,-0.0] | 0.049* | 0 | [-0.05,0.04] | 0.902 |
| Frontal pole | -0.08 | [-0.12,-0.04] | <0.001***a | -0.02 | [-0.06,0.03] | 0.465 |
| Fusiform | -0.08 | [-0.11,-0.04] | <0.001***a | -0.05 | [-0.09,0.0] | 0.051 |
| Inferior parietal | -0.01 | [-0.04,0.02] | 0.623 | -0.06 | [-0.1,-0.01] | 0.011*a |
| Inferior temporal | -0.06 | [-0.09,-0.02] | 0.002**a | -0.04 | [-0.09,0.0] | 0.052 |
| Insula | -0.03 | [-0.06,0.01] | 0.188 | -0.01 | [-0.06,0.04] | 0.644 |
| Isthmus cingulate | -0.06 | [-0.1,-0.02] | 0.002**a | -0.03 | [-0.08,0.02] | 0.258 |
| Lateral occipital | -0.02 | [-0.06,0.01] | 0.126 | -0.11 | [-0.15,-0.07] | <0.001***a |
| Lateral orbitofrontal | -0.13 | [-0.17,-0.1] | <0.001***a | -0.04 | [-0.08,0.01] | 0.133 |
| Lingual | -0.06 | [-0.09,-0.02] | 0.003**a | -0.12 | [-0.17,-0.08] | <0.001***a |
| Medial orbitofrontal | -0.13 | [-0.17,-0.09] | <0.001***a | 0 | [-0.04,0.05] | 0.939 |
| Middle frontal (Rostral) | -0.12 | [-0.15,-0.08] | <0.001***a | -0.02 | [-0.06,0.03] | 0.429 |
| Middle temporal | -0.06 | [-0.1,-0.03] | <0.001***a | -0.05 | [-0.09,-0.01] | 0.022* |
| Paracentral | -0.08 | [-0.11,-0.04] | <0.001***a | -0.07 | [-0.11,-0.02] | 0.004**a |
| Parahippocampus | -0.02 | [-0.05,0.02] | 0.459 | -0.04 | [-0.09,0.0] | 0.069 |
| Parsopercularis | -0.03 | [-0.07,0.0] | 0.074 | -0.01 | [-0.06,0.03] | 0.538 |
| Parsorbitalis | -0.09 | [-0.13,-0.05] | <0.001***a | -0.03 | [-0.07,0.02] | 0.268 |
| Parstriangularis | -0.08 | [-0.11,-0.04] | <0.001***a | 0 | [-0.05,0.04] | 0.82 |
| Pericalcarine | -0.01 | [-0.04,0.03] | 0.717 | -0.09 | [-0.13,-0.04] | <0.001***a |
| Postcentral | -0.01 | [-0.05,0.02] | 0.449 | -0.1 | [-0.14,-0.06] | <0.001***a |
| Posterior cingulate | -0.04 | [-0.08,-0.0] | 0.027* | 0 | [-0.05,0.05] | 0.992 |
| Precentral | -0.02 | [-0.05,0.02] | 0.267 | -0.06 | [-0.1,-0.01] | 0.013*a |
| Precuneus | -0.06 | [-0.09,-0.02] | 0.004**a | -0.05 | [-0.1,-0.0] | 0.035* |
| Superior frontal | -0.1 | [-0.13,-0.06] | <0.001***a | -0.01 | [-0.06,0.03] | 0.633 |
| Superior parietal | -0.02 | [-0.06,0.01] | 0.183 | -0.07 | [-0.11,-0.02] | 0.003**a |
| Superior temporal | -0.05 | [-0.09,-0.02] | 0.004**a | -0.04 | [-0.08,0.01] | 0.103 |
| Superior temporal (Banks) | -0.01 | [-0.05,0.03] | 0.664 | -0.06 | [-0.11,-0.01] | 0.012*a |
| Supramarginal | 0.01 | [-0.02,0.04] | 0.625 | -0.03 | [-0.07,0.0] | 0.084 |
| Temporal pole | -0.06 | [-0.1,-0.02] | 0.003**a | 0 | [-0.05,0.04] | 0.941 |
| Transverse temporal | -0.03 | [-0.07,0.01] | 0.108 | -0.08 | [-0.13,-0.04] | <0.001***a |

**Table 3.** Parameter estimates from the neuroinflammation theory of overeating looking at the associations between ADI and BMI at 9/10-years-old and subcortical volume at 11/12-years-old. Associations were tested using linear random mixed effects covaried for puberty, sex, Race/ethnicity and education with a random effect of scanner type was utilized to identify ROIs at 11/12-years-old that were significantly associated with ADI and BMI at 9/10-years-old. Significant effects were corrected separated by modality (e.g., cortical thickness, volume) and model (e.g., ADI, BMI). ^a^survived multiple correction testing using the Benjamini-Hochberg method. Salmon-colored rows highlight the ROIs that showed significant associations with BMI and ADI. Blue rows highlight regions that were significant but there was no overlap between BMI and ADI. Gray rows had no significant associations.

| Neuroinflammation theory of overeating |  |  |  |  |  |  |
| --- | --- | --- | --- | --- | --- | --- |
| ROI (11-12-yrs) | Associations with BMI at 9-10-yrs |  |  | Associations with ADI at 9-10-yrs |  |  |
|  | Beta | 95%CI | *p* | Beta | 95%CI | *p* |
| Accumbens area | -0.02 | [-0.05,0.02] | 0.365 | -0.09 | [-0.14,-0.05] | <0.001***<sup>a</sup> |
| Amygdala | -0.03 | [-0.06,0.01] | 0.11 | -0.11 | [-0.15,-0.07] | <0.001***<sup>a</sup> |
| Caudate | 0.01 | [-0.02,0.05] | 0.467 | -0.13 | [-0.18,-0.08] | <0.001***<sup>a</sup> |
| Hippocampus | 0 | [-0.03,0.04] | 0.922 | -0.12 | [-0.16,-0.07] | <0.001***<sup>a</sup> |
| Pallidum | 0.04 | [0.0,0.08] | 0.028* | -0.09 | [-0.13,-0.04] | <0.001***<sup>a</sup> |
| Putamen | -0.03 | [-0.06,0.01] | 0.118 | -0.13 | [-0.17,-0.08] | <0.001***<sup>a</sup> |
| Thalamus | 0.03 | [-0.0,0.07] | 0.067 | -0.14 | [-0.18,-0.09] | <0.001***<sup>a</sup> |
| Ventral DC | 0.07 | [0.04,0.11] | <0.001***<sup>a</sup> | -0.1 | [-0.14,-0.06] | <0.001***<sup>a</sup> |

### Neural stress theory of overeating

Brain regions at 9/10-years-old that were associated with ADI at 9/10-years-old and BMI at 11/12-years-old are presented in **Tables 4–5**. Results from the parallel mediation model indicated that brain structure at 9/10-years-old partially mediated the association between ADI at 9/10-years-old on BMI at 11/12-years-old, which suggests that these associations may be drive by stress-related effects of living in an area deprived of resources. Positive and negative brain structure mediators are displayed in **Fig4A**. ADI at 9/10-years-old was positively associated with BMI at 11/12-years-old (C path, Total direct effect, **Fig4B**) but negatively associated with cortical thinning of the lateral orbitofrontal cortex and lingual gyrus and decreased subcortical volume of the Ventral DC at 9/10-years-old (*a* path, **Fig4B**). Cortical thinning and larger subcortical volume of the Ventral DC at 9/10-years-old were associated with greater BMI at 11/12-years-old (*b* path, **Fig4B**). No other regions that were associated with ADI at 9/10-years-old and BMI at 11/12-years-old were significant mediators.

**Fig4.**
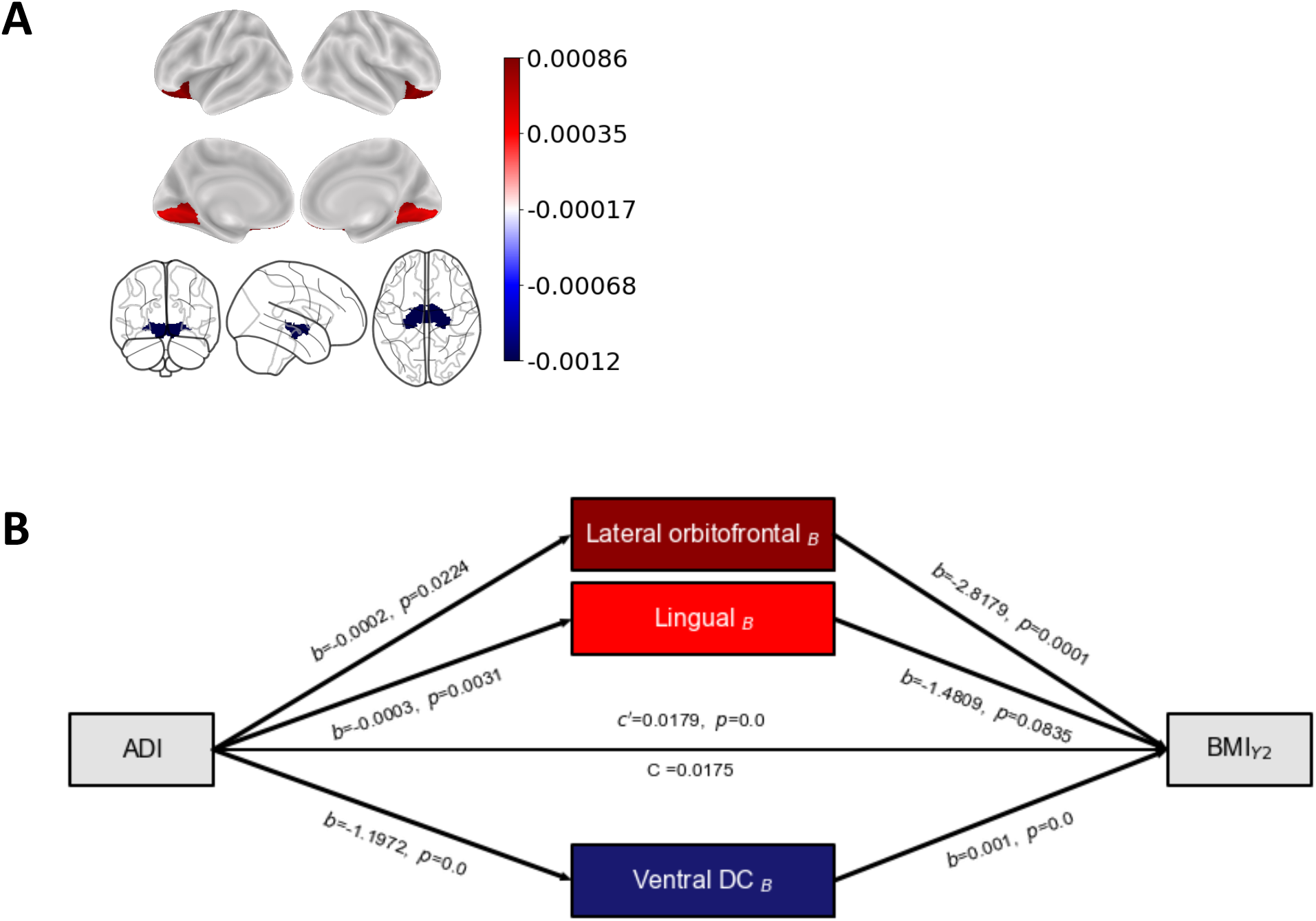
Testing the causal pathway of the neuronal stress theory of overeating. Mediating effects of brain structure in the lateral orbitofrontal cortex, lingual gyrus, and Ventral DC at 9/10-years-old on ADI and brain structure at 11/12-years-old. **A)** Visual representation of the brain regions that showed mediated the effects of ADI on BMI at 11-to-12-years colored by the strength of the indirect association. **B)** Mediation models where the colored boxes reflect the strength (and direction) of the indirect effects. B=baseline (aged 9/10-years-old). Y2=two-year follow-up (aged 11/12-years-old). ADI = area deprivation index. BMI = body mass index. Total effects are represented by *c’*, direct effects of ADI are represented by C while *a* and *b* values refer to the association of ADI on BMI and BMI on brain structure, respectively. All *a, b, c*, and *c’* values are unstandardized regression coefficients. Significance testing was carried out by bias-corrected bootstrapping (n=10,000) 95% confidence intervals.

**Table 4.**
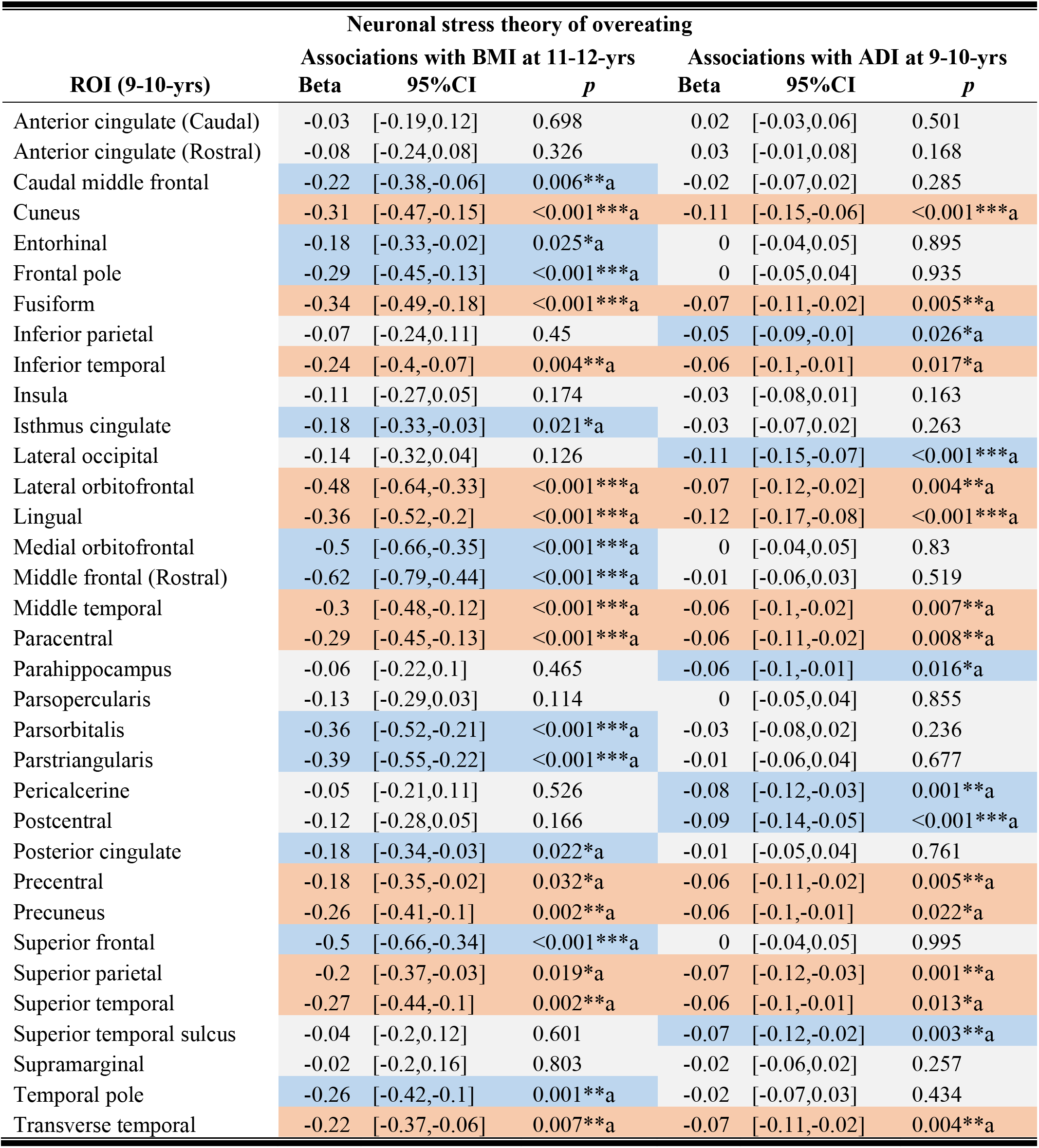
Parameter estimates from the neuronal stress theory of overeating assessing the associations between ADI and cortical thickness at 9/10-years-old and BMI at 11-to-12-year-old. Linear random mixed effects covaried for puberty, sex, race/ethnicity and education, scanner type (random effect) to identify ROIs at 9/10-years-old that were significantly associated with ADI at 9/10-years-old and BMI at 11/12-years-old. Correction was conducted separately by model (e.g., ADI, BMI). ^a^survived multiple correction via the Benjamini-Hochberg method. Salmoncolored rows highlight the ROIs that were significantly associated with BMI and ADI. Blue rows highlight regions that were significant but there was no overlap between BMI and ADI. Gray rows had no significant associations.

**Table 5.**
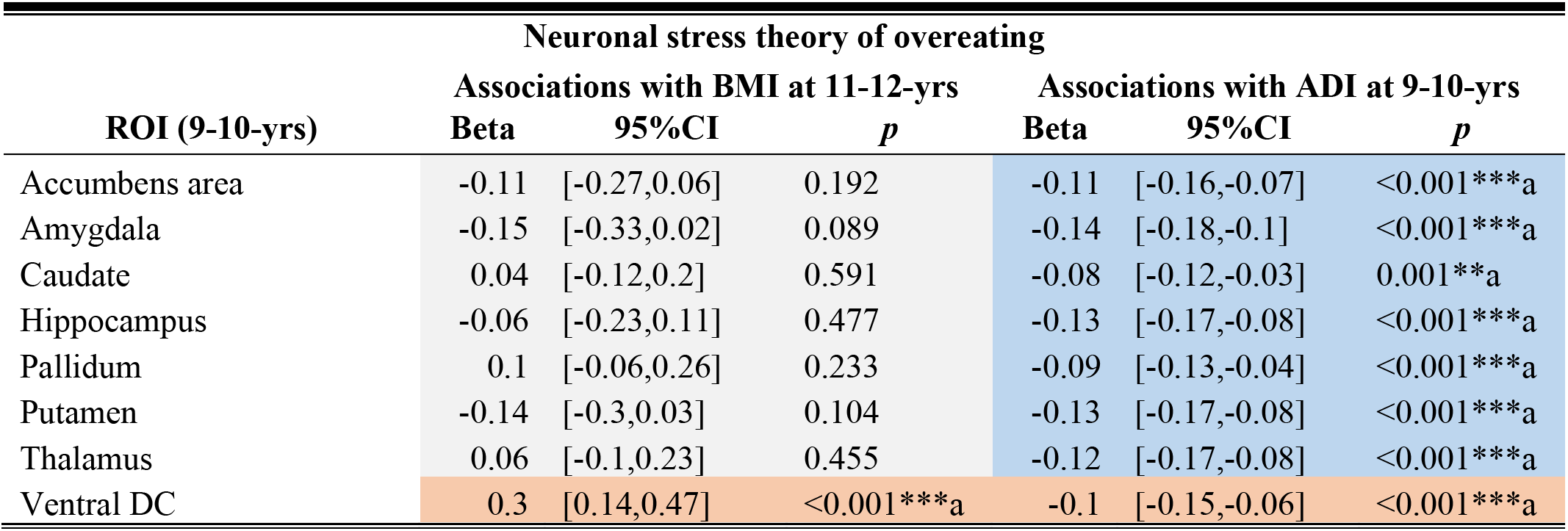
Parameter estimates from the neuronal theory of overeating looking at the associations between ADI and subcortical volume at 9/10-years-old and BMI at 11-to-12-year-old. Associations were tested using linear random mixed effects covaried for puberty, sex, Race/ethnicity and education with a random effect of scanner type was utilized to identify ROIs at 9/10-years-old that were significantly associated with ADI at 9/10-years-old and BMI at 11/12-years-old. Significant effects were corrected separated by modality (e.g., cortical thickness, volume) and model (e.g., ADI, BMI). ^a^survived multiple correction testing using the Benjamini-Hochberg method. Salmon-colored rows highlight the ROIs that showed significant associations with BMI and ADI. Blue rows highlight regions that were significant but there was no overlap between BMI and ADI. Gray rows had no significant associations.

| Neuronal stress theory of overeating |  |  |  |  |  |  |
| --- | --- | --- | --- | --- | --- | --- |
| ROI (9-10-yrs) | Associations with BMI at 11-12-yrs |  |  | Associations with ADI at 9-10-yrs |  |  |
|  | Beta | 95%CI | *p* | Beta | 95%CI | *p* |
| Accumbens area | -0.11 | [-0.27,0.06] | 0.192 | -0.11 | [-0.16,-0.07] | <0.001***<sup>a</sup> |
| Amygdala | -0.15 | [-0.33,0.02] | 0.089 | -0.14 | [-0.18,-0.1] | <0.001***<sup>a</sup> |
| Caudate | 0.04 | [-0.12,0.2] | 0.591 | -0.08 | [-0.12,-0.03] | 0.001**<sup>a</sup> |
| Hippocampus | -0.06 | [-0.23,0.11] | 0.477 | -0.13 | [-0.17,-0.08] | <0.001***<sup>a</sup> |
| Pallidum | 0.1 | [-0.06,0.26] | 0.233 | -0.09 | [-0.13,-0.04] | <0.001***<sup>a</sup> |
| Putamen | -0.14 | [-0.3,0.03] | 0.104 | -0.13 | [-0.17,-0.08] | <0.001***<sup>a</sup> |
| Thalamus | 0.06 | [-0.1,0.23] | 0.455 | -0.12 | [-0.17,-0.08] | <0.001***<sup>a</sup> |
| Ventral DC | 0.3 | [0.14,0.47] | <0.001***<sup>a</sup> | -0.1 | [-0.15,-0.06] | <0.001***<sup>a</sup> |

## Discussion

The current study was the first to evaluate two competing (and complimentary) theories of how ADI may relate to childhood obesity and brain development during a two-year period in adolescence. Here, we provide support for the neuroinflammation theory of overeating which posits that the relationship between ADI and BMI at 9/10-years-old and brain structure at 11/12-years-old may be influenced by a lack of access to healthy food, and that the neuroinflammatory effects of unhealthy food may alter brain structure later in development (Guillemot-Legris & Muccioli, 2017; Mullins et al., 2020). We also found evidence to support the neuronal stress theory of overeating, which posited that the relationship between brain structure and ADI 9/10-years-old and BMI at 11/12-years-old may be influenced by the stressful impacts of living in a neighborhood with fewer resources, which may trigger stress-induced overeating. Together, the overall pattern of findings highlights the implications of greater area deprivation for potential child health disparities in obesity and alterations in brain structure. Greater area deprivation is a proxy for fewer neighborhood and environmental resources that are a result of systemic inequities. Therefore, these findings add to our understanding of how the social determinants of health may influence childhood obesity (Dixon et al., 2021). To our knowledge, this is the first study to test two parallel and competing theoretical models explaining relationships between area deprivation, and brain development and childhood obesity.

The Ventral diencephalon (Ventral DC) emerged as an important region that may explain how area deprivation is associated with both childhood obesity and brain structure from a neuroinflammation and neuronal-stress framework. The Ventral DC is an area that incorporates grey and white matter structures, such as the hypothalamus, mamillary body, medial geniculate nucleus, red nucleus, and parts of the basal ganglia (e.g., nigra and subthalamic nuclei) (Herrero et al., 2002); these regions are involved in food intake signaling and dopamine secretion (Gantz et al., 2018). The neuroinflammation theory of overeating suggested that greater ADI may directly influence Ventral DC volume at 11/12-years-old from the indirect associations on BMI at 9/10-years-old. From a neuroinflammatory perspective, overconsumption of unhealthy foods may alter food intake neurons and, in the hypothalamus, cause overexpression of dopamine (a reward response neurotransmitter) (Mullins et al., 2020); overexpression of dopamine is associated with hypersensitivity to rewards (Kroemer & Small, 2016; Stice & Yokum, 2016), which is thought to promote overeating in youth (Adise et al., 2018). Another explanation could be that larger volumes of the Ventral DC at 11/12-years-old occur due to an adaptive response based on deficits in other brain regions (de Groot et al., 2017), which may also be affected by neuroinflammation (Castanon et al., 2014), such as cortical thinning of the cuneus, lingual, and paracentral gyrus. The cuneus and lingual gyrus are associated with visual processing, and altered functional responses in these regions to food cues have been thought to be indicative of increased attentional bias to food (Gearhardt et al., 2014; Goldschmidt et al., 2018).

We also found support for the neuronal stress theory of overeating, as, at 9/10-years-old, greater ADI was associated with reduced subcortical volume of the Ventral DC *and* cortical thinning in regions associated with reward processing (e.g., lateral orbitofrontal cortex and attentional biases to food (e.g., lingual gyrus) (Gearhardt et al., 2014; Rolls, 2015). These structural variations were associated with increased BMI at 11/12-years-old. These findings confer with the literature suggesting that area deprivation is associated with smaller volumes and thinner cortex (Brito & Noble, 2014; Noble et al., 2015), possibly due to neuronal atrophy from overstimulation of the hypothalamic response to the chronic stress of living in an area deprived of resources (Miller & Spencer, 2014). Smaller volumes are thought to decrease dopamine availability (Mullins et al., 2020), which may trigger overeating as an attempt to (1) normalize reward processing (Stice & Yokum, 2016), or (2) downregulate stress (Razzoli & Bartolomucci, 2016), or (3) reduced dopamine availability may also increase BMI by reducing spontaneous exercise (Friend et al., 2017). However, the ABCD Study^®^ did not assess dopamine expression, so future research is needed to evaluate this hypothesis in youth. It is also important to note that within the neuronal stress theory of overeating, we observed a positive correlation between Ventral DC volume and BMI at 11/12-years-old, suggesting that the relationship between ADI and brain structure at 9/10-years-old and BMI at 11/12-years-old is complex. One reason to explain this may be related to the bidirectional effects of stress-induced overeating: Animal models show that chronic stress triggers the hypothalamus to increase food intake, but, because stress also increases energy expenditure, stress-induced overeating does not always lead to weight gain (Razzoli & Bartolomucci, 2016). Although these findings do support that area deprivation may relate to brain structure and BMI possibly due to the stressful impacts of living in a deprived neighborhood, future research is needed to tease apart these mechanisms.

Interestingly, brain regions associated with inhibitory control showed no effects of ADI or BMI for either theory to explain the associations of area deprivation, brain development, and obesity. This is contrary to another ABCD Study^®^, in which ADI moderated the strength of the relationship between BMI at 9/10-years-old and cortical thinning at 9/10-years-old in inhibitory control regions such as the inferior frontal and medial frontal cortex (Hall et al., 2021). However, cross-sectional studies provide little insight into temporal stability and causal inferences, particularly in a sample that is undergoing brain maturation in reward and inhibitory control regions (Shulman et al., 2016). Our present findings may be indicative of a developmental effect (Shaw et al., 2006), in which area deprivation accelerates maturation of brain regions associated with reward processing occur faster than those involved in inhibitory control (Luna et al., 2004; Mills et al., 2014). Thus, within this age range, greater area deprivation may accelerate reward sensitivity that leads to increased food intake while an immature inhibitory control system lacks the maturity to suppress urges to overeat. Accelerated development of reward sensitivity and food intake signaling may have downstream effects on cognition, and future releases of the ABCD Study^®^ will provide insight into how ADI and the brain change in respect to obesity over time, particularly within this developmental window.

Our findings reinforce how much the human-made landscape of the communities in which we live influence child health beyond individual sociodemographics and economics. Importantly, the relationships between area deprivation, brain structure, and BMI were independent of both sociodemographic (e.g., race) *and* socioeconomic (e.g., education) factors. As ADI takes into account several aspects of social and environmental factors, it may be a better measure to capture how social determinants of health relate to poorer health outcomes, like childhood obesity. Therefore, our results serve as a reference to study the longitudinal associations of the brain and obesity development over time.

### Strengths and limitations

The findings in this manuscript offer a unique contribution to the literature by demonstrating how lack of resources in the environment can differentially affect obesity via two different causal frameworks. However, there were a few limitations. Our methodological framework was setup in a temporal fashion to test causal pathways. Yet, caution should still be exercised when interpreting results. The first MRI available was at 9/10 years old, so it is possible that other pre-existing conditions may explain these relationships. The ABCD Study^®^ did not collect markers of oxidative stress nor inflammation, therefore we are limited in regard to the inferences that can be made. In addition, other potential mediators that were not assessed or measured may partially or equally explain our results, such as gene by environment interactions. It is also not known how these observations may change after pubertal onset and with normative brain maturation that occurs over the course of development. Related to this, we were not able to assess longitudinal change within these causal frameworks as additional data collections are needed in order to have temporal specificity and longitudinal change in the same model. However, future releases of MRI data will allow for these investigations. Although ADI is a better measure of deprivation than family income or parental education, it may be subject to some biases. For example, environmental stressors in urban versus rural areas may be different, and it is hard to tease apart these differences within ADI. Additional environmental data, such as access to grocery stores and objective cortisol measures may help to clarify some of the finegrained associations observed in this study. Future releases of the ABCD Study^®^ may be able to offer insight into more fine-grained mechanisms, but this current study serves as a reference point.

## Conclusions

These findings highlight the crucial role of area deprivation for obesity and brain development during pre- and early adolescence. Our findings show that that ADI may relate to health and development from a neuroinflammatory and neuronal stress framework. Without proper intervention from a public health policy perspective, our findings support the notion that communities that have been disproportionately affected by greater neighborhood deprivation will continue to be at risk for deleterious health outcomes associated with a lack of environmental resources. This dovetails previous research stating how important neighborhood resources are for optimal health. The neighborhood environment factor that can be modified by improving resource access (e.g., adding health food options, increasing green space, creating safe environments, increasing access to affordable health care), which can greatly influence health outcomes. Thus, public health policy may wish to advocate for more funds to improve resources in deprived areas as a means to decrease negative health outcomes.

## Funding acknowledgements

Data used in the preparation of this article were obtained from the Adolescent Brain Cognitive Development™ Study^®^ (https://abcdstudy.org/), held in the NIMH Data Archive (NDA). The **ABCD Study^®^** is supported by the National Institutes of Health and National Institute on Drug Abuse and additional federal partners under award numbers U01DA041022, U01DA041028, U01DA041048, U01DA041089, U01DA041106, U01DA041117, U01DA041120, U01DA041134, U01DA041148, U01DA041156, U01DA041174, U24DA041123, and U24DA041147. A full list of supporters is available at https://abcdstudy.org/federal-partners/. A listing of participating sites and a complete listing of the study investigators can be found at https://abcdstudy.org/principal-investigators/. The **ABCD Study^®^** consortium investigators designed and implemented the study and/or provided data but did not necessarily participate in analysis or writing of this report. This manuscript reflects the views of the authors and may not reflect the opinions or views of the NIH or other **ABCD Study^®^** consortium investigators. The **ABCD Study^®^** data repository grows and changes over time. The **ABCD Study^®^** data used in this report came from https://doi.org/10.15154/1503209.

