## Supplemental Materials for "Relating neighborhood deprivation to childhood obesity in the ABCD Study®: evidence for theories of neuroinflammation and neuronal stress"

**Demographic assessments**. Caregiver report of the child’s race/ethnicity, date of birth, and sex at birth were obtained at the baseline visit. The child’s age at the time of each visit was recorded in months. Socioeconomic status was operationalized using the highest education of the household, of which there were 29 categories. Education was collapsed into five groups: <High school (HS), HS/Generalized Education Diploma (GED), Some college, Four-year degree (Bachelor’s degree), Postgraduate education. Race and ethnicity had 22 options, which were collapsed into five groups: White, Black, Hispanic, Asian, and Mixed/Other.

**Behavioral Inhibition System / Behavioral Approach System (BIS/BAS)***:* Youth completed a modified and original BIS/BAS questionnaire^1^, which is commonly used to assess trait-based reward and inhibitory control. The questionnaire consists of 24 questions that are summarized into three BAS subscales and one BIS scale. The behavioral approach system (BAS) assesses goal-directed behaviors with subscales for reward responsiveness, fun seeking, and drive. The behavioral inhibition system (BIS) assesses avoidance-type behaviors and only has one subscale. Questions for each subscale are scored on a 4-point scale from 1 (very true) to 4 (very false). Answers on the modified and original questionnaires were highly correlated (*r*=0.9), so only the modified BIS/BAS was used in the analysis. Descriptions of each measure are presented in the **Supplemental Materials Table S3**.

**Urgency, Premeditation, Perseverance, Sensation Seeking, and Positive Urgency (UPPS-P):** The UPPS-P is a 59-item questionnaire that was answered by youth. The five subscales identify traits as outlined in the scale title. Questions for each subscale are scored on a 4-point scale from 1 (agree strongly) to 4 (disagree strongly). Descriptions of each measure are presented in the **Supplemental Materials Table S3**.

**Elastic net regression:**

Initially, the entire dataset was randomly split into an 80% training dataset for model construction and a 20% testing set to access model generalizability. Within the training dataset, separate models were trained and evaluated with 5-fold cross-validation (CV), stratified to preserve the ratio of WS_HW_ to WG participants within each fold. A measure of generalizability was accessed by applying the highest performing of the five models, as estimated by performance on their respective validation folds, to the previously unseen testing set. We report performance using the binary classification metric area under the curve (AUC), where AUC_train_ refers to AUC on the model’s associated validation fold in the training set and AUC_test_ to AUC as calculated from the testing set. The implementation of the logistic elastic-net regression model included a further nested hyper-parameter search across 60 random combinations of values for both strength of regularization as well as ratio between L1 and L2 regularization. All machine learning models, training, and evaluation were conducted in Python through the Brain Predictability Toolbox package^2^. Brain features made available to the elastic-net model consisted of cortical Destrieux (n=148) and subcortical ROIs (n=14) for cortical thickness, surface area, subcortical volume, DTI FA, and DTI MD estimates (n_total_=648), as well as intracranial volume and motion estimates for the DTI scan. A set of nonbrain features were also made available as input features, which consisted of sex, age, puberty, highest household education (a proxy for socioeconomic status), race/ethnicity, intracranial volume, motion estimates for the DTI scan, and scanner ID. As a quality control step, the top and bottom 1% of the data for each ROI were winsorized (i.e., values <1% and >99% percentiles, respectively, were set to a truncated value) and then further normalized (each feature scaled to mean=0 and standard deviation=1 within that training sample). Both procedures were performed in a properly nested manner, where estimates of both 1^st^ and 99^th^ quantiles as well as other sample dependent measures were estimated only within training data folds. These analyses were conducted on 747/748 youth because one subject was missing fractional anisotropy estimates for two ROIs. Although the ABCD **Study^®^** often enrolled siblings, in this subset of the data, 98% (n=731) of youth were singletons. However, after outlier exclusion, the test set had only 1 sibling pair within the WS_HW_ group, and, thus, independence issues in the test set were not of concern.

**Weight gain model confirmation:** Previously, one-year weight gain was found to not be confounded by BMI at baseline^3^. Again, to make sure weight status was not confounding results, we removed WG youth who were classified as overweight or obese at baseline and reran the elastic-net regression. To account for a decrease in sample size, we increased the number of folds 10, otherwise, the parameters remained the same.

**Table S1.** The number of subjects available based on each exclusion criterion applied.

|  | | | **n** |
| --- | --- | --- | --- |
| Y2 released data | | | 6571 |
| Not underweight (i.e., BMI %ile >5) | | | 5882 |
| No medications known to affect food intake | | | 5882 |
| No learning disabilities or psychiatric disorders | | | 5882 |
| No eating disorders based on caregiver report on the KSADS | | | 4941 |
| Complete data for sex, age, puberty, race, education | | | 4709 |
| No height measurement error (i.e., decrease in height between visits) | | | 4617 |
| Met WS_HW_/WG criteria | | | 1034 |
|  | WS_HW_ (n=709) | WG (n = 316) | |
| Passed FreeSurfer QC | 587 | 256 | |
| Acceptable T_1_ | 584 | 254 | |
| Acceptable T_2_ | 544 | 238 | |
| Acceptable DTI | 529 | 226 | |
| No missing tabulated data | 527 | 221 | |
| Final sample | 527 | 221 | |

*Note.* Y2 = year 2; BMI %ile = Body mass index percentile according to the CDC sex-age-height-weight-specific growth curves for youth; WS_HW_ = Healthy weight, Weight Stable; WG = Weight Gain; QC = quality control; T_1_ = T_1_-weighted image; T_2_ = T_2_-weighted image; DTI = diffusion tensor imaging. KSADS = Kiddie schedule for affective disorders and schizophrenia for school-age youth.

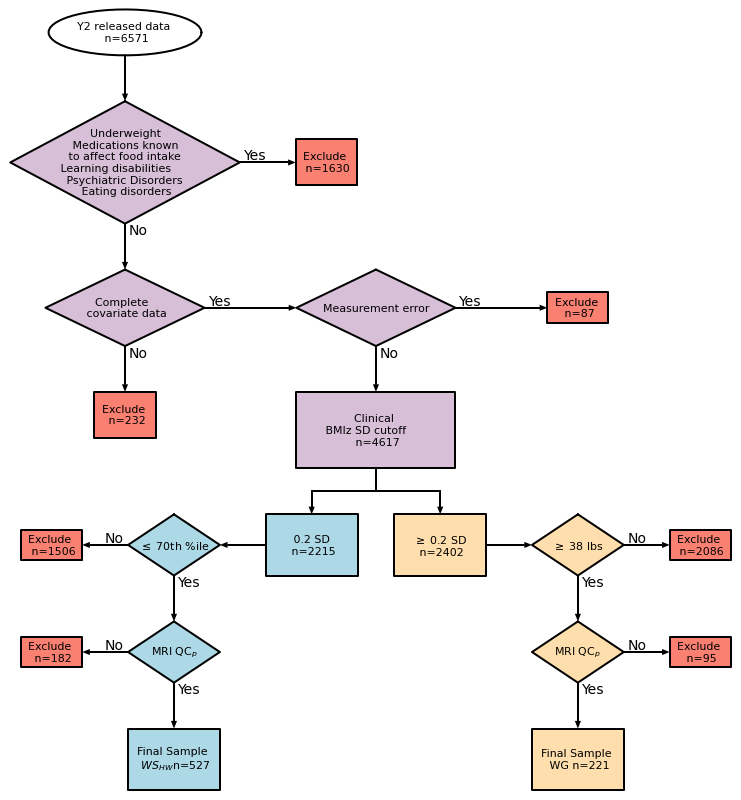

**Figure S1**: An overview of group selection. Y2= year 2; BMIz = BMI z-score; SD = standard deviation; %ile = BMI percentile; MRI = Magnetic Resonance Imaging; MRI QC*_p_* = passed MRI quality control parameters; WG = weight gain; WS_HW_ = Healthy Weight, Weight Stable.

**Table S2.** The number of youth in the health weight, Weight Stable (WS_HS_) and Weight Gain (WG) groups as defined in Adise et al., (2021) ^19^ that met criteria for WS and WG by the criteria outlined in this manuscript

|  | WS_HW_ at Y1 (n=637) | WG at Y1 (n=172) |
| --- | --- | --- |
| Had BMI data at Y2 | 617 | 165 |
| No missing covariates at any time point | 615 | 163 |
| No eating disorders at Y1 or Y2 | 536 | 157 |
| No medications known to affect food intake at Y1 or Y2 | 536 | 157 |
| Had MRI data at Y2 | 475 | 140 |
| Passed FreeSurfer QC at Y2 | 473 | 138 |
| Passed T_1_ MRI QC at Y2 | 473 | 137 |
| Passed T_2_ MRI QC at Y2 | 460 | 127 |
| Passed DTI QC at Y2 | 456 | 127 |
| Had BMI data at Y2 | 452 | 126 |
| No BMI measurement error | 448 | 126 |
| Met BMIz SD criteria from baseline to Y2 | 277 | 113 |
| Met WS_HW_/WG criteria | 249 | 58 |
| No missing tabulated MRI data | 249 | 56 |

*Note*. Weight criteria for youth in the WS_HW_ group corresponded to a BMI percentile < 70% at all time points and a BMIz SD <0.2 (clinical cut off). Weight criteria for youth in the WG group corresponded to weight gain > 38 lbs. across all time points in addition to a BMIz SD >=0.2 (clinical cutoff). BMI = Body Mass Index. BMIz = BMI z-score; T_1_ = T_1_-weighted image; T_2_ = T_2_-weighted image; DTI = diffusion tensor imaging; Y1 = year 1; Y2 = year 2. MRI = Magnetic resonance imaging; QC = quality control; STD = standard deviation.

**Table S3.** Variable descriptions.

| **Variable Groupings and Names in Data Release** | **Variable Descriptions** |
| --- | --- |
| **Kiddy schedule for affective disorders and schizophrenia (KSADS)** |  |
| Binge eating disorder | Eating much more than normal in a short period of time, eating when not hungry. |
| Anorexia nervosa | Restrictive eating, underweight (BMI<18.5 or 5^th^ %ile) |
| Bulimia nervosa | Binging (i.e., eating a lot) and purging (i.e., getting rid of) behavior |
| Eating disorder not otherwise specified (EDOS) or Other Specified Feeding or Eating Disorder (OSFED) | Captures individuals who do not meet diagnostic criteria for anorexia nervosa or bulimia nervosa but still present with eating disorder-like symptoms |
| **Behavioral Inhibition System/ Behavioral Approach System** |  |
| Reward responsiveness | Sensitivity to pleasant reinforcers in the environment |
| Fun seeking | Motivation to find novel rewards spontaneously |
| Drive | Motivation to follow goals |
| Inhibition | Inhibitory control |
| **Urgency, Premeditation, Perseverance, Sensation Seeking, and Positive Urgency (UPPS-P)** |  |
| Negative urgency | The tendency to act rashly (i.e., impulsively) under an intense negative mood |
| Positive urgency | The tendency to act rashly (i.e., impulsively) while under an intense positive mood |
| Sensation seeking | The tendency to seek out novel and thrilling experiences |
| Lack of planning/premeditation | The tendency to not take into account the consequences of actions |
| Lack of perseverance | The tendency to quit a task when it becomes difficult, long, or boring. |

**Table S4.** Prenatal characteristics for youth in the Healthy Weight, Weight Stable (WS_HW_) and Weight Gain (WG) groups and the rest of the sample with available data for year 2 (Other).

|  | **WS_HW_ (n=527)** | **WG (n=225)** | ***p _group_*** | **Other** | ***p _all_*** |
| --- | --- | --- | --- | --- | --- |
| Birth Weight [*M* (*SD*)] | 13.9 (4.5) | 14.1 (4.4) | 0.6 | 13.7 (4.4) | 0.3 |
| Born Premature [*n* (%)] |  |  |  |  |  |
| No | 415 (78.9) | 176 (79.6) | 0.6 | 3070 (79.3) | 0.6 |
| Yes | 106 (20.2) | 41 (18.6) |  | 768 (19.9) | 0.5 |
| Refused to Answer | 5 (1.0) | 4 (1.8) |  | 31 (0.8) |  |
| Weeks Premature [*M* (*SD*)] | 4.6 (2.2) | 5.3 (2.1) | 0.7 | 15.3 (101.0) |  |
| Prenatal tobacco (short) exposure [*n* (%)] |  |  |  |  |  |
| No | 467 (88.6) | 173 (78.3) | 0.001 | 3343 (86.4) | 0.005 |
| Yes | 52 (9.9) | 40 (18.1) |  | 452 (11.7) |  |
| Refused to Answer | 8 (1.5) | 8 (3.6) |  | 73 (1.9) |  |
| Prenatal tobacco (continued) exposure [*n* (%)] |  |  |  |  |  |
| No | 498 (94.5) | 198 (89.6) | 0.04 | 2679 (69.3) | 0.7 |
| Yes | 22 (4.2) | 15 (6.8) |  | 973 (25.2) |  |
| Refused to Answer | 7 (1.3) | 8 (3.6) |  | 216 (5.6) |  |
| Prenatal alcohol (short) exposure [*n* (%)] |  |  |  |  |  |
| No | 356 (67.6) | 152 (68.8) | 0.8 | 3656 (94.5) | 0.03 |
| Yes | 133 (25.2) | 56 (25.3) |  | 151 (3.9) |  |
| Refused to Answer | 38 (7.2) | 13 (5.9) |  | 61 (1.6) |  |
| Prenatal alcohol (continued) exposure [*n* (%)] |  |  |  |  |  |
| No | 506 (96.0) | 208 (94.1) | 0.03 | 3710 (95.9) | 0.09 |
| Yes | 16 (3.0) | 5 (2.3) |  | 100 (2.6) |  |
| Refused to Answer | 5 (0.9) | 8 (3.6) |  | 58 (1.5) |  |

*Note. p_group_* = significant group differences between WG and WS. *p_all_* = significant differences between WS_HW_, WG, and the rest of the sample. *M* = mean; *SD* = standard deviation; *n* = sample size; % = Percent. Short = substance use discontinued after pregnancy knowledge; continued = substance use continued throughout pregnancy. *p*-values represent significance testing for *t*-tests and chi-squared testing.

**Table S5.** Results from the mixed effects model examining how brain regions that were predictive of one-year weight gain ^3^ changed in response to two-years of weight gain.

| ROI |  | *F* | *p* |
| --- | --- | --- | --- |
| *Cortical Thickness* |  |  |  |
| Anterior transverse collateral sulcus LH |  |  |  |
|  | Group | 0.61 | 0.433 |
|  | Time | 0.16 | 0.685 |
|  | Group x Time | 0.13 | 0.718 |
| Frontomarginal gyrus and sulcus RH |  |  |  |
|  | Group | 19.2 | < 0.001 ^a^ *** |
|  | Time | 2.18 | 0.140 |
|  | Group x Time | 9.86 | < 0.001 ^a^ *** |
| Pericallosal sulcus LH |  |  |  |
|  | Group | 0.52 | 0.467 |
|  | Time | 0.08 | 0.774 |
|  | Group x Time | 0.97 | 0.322 |
| Rectal gyrus LH |  |  |  |
|  | Group | 1.87 | 0.171 |
|  | Time | 0.07 | 0.783 |
|  | Group x Time | 0.31 | 0.574 |
| Sulcus intermedius primus (of Jensen) LH |  |  |  |
|  | Group | 2.25 | 0.133 |
|  | Time | 3.10 | 0.078 |
|  | Group x Time | 2.53 | 0.111 |
| *Surface Area* |  |  |  |
| Angular gyrus RH |  |  |  |
|  | Group | 0.00 | 0.983 |
|  | Time | 0.58 | 0.444 |
|  | Group x Time | 0.98 | 0.321 |
| Anterior occipital sulcus RH |  |  |  |
|  | Group | 0.15 | 0.698 |
|  | Time | 1.53 | 0.215 |
|  | Group x Time | 0.92 | 0.335 |
| Anterior transverse temporal gyrus LH |  |  |  |
|  | Group | 0.00 | 0.969 |
|  | Time | 0.12 | 0.722 |
|  | Group x Time | 4.02 | 0.045 * |
| Parahippocampal gyrus LH |  |  |  |
|  | Group | 2.74 | 0.098 |
|  | Time | 0.01 | 0.911 |
|  | Group x Time | 0.78 | 0.374 |
| Superior segment of the circular sulcus of the insula RH |  |  |  |
|  | Group | 0.29 | 0.586 |
|  | Time | 0.00 | 0.987 |
|  | Group x Time | 0.05 | 0.809 |
| *Fractional Anisotropy* |  |  | . |
| Intraparietal sulcus and transverse parietal sulci RH |  |  |  |
|  | Group | 1.36 | 0.243 |
|  | Time | 2.34 | 0.125 |
|  | Group x Time | 2.30 | 0.129 |
| Lateral orbital sulcus LH |  |  |  |
|  | Group | 0.11 | 0.739 |
|  | Time | 1.55 | 0.212 |
|  | Group x Time | 0.00 | 0.977 |
| Paracentral gyrus and sulcus LH |  |  |  |
|  | Group | 0.58 | 0.445 |
|  | Time | 3.25 | 0.071 |
|  | Group x Time | 0.19 | 0.662 |
| *Mean Diffusivity* |  |  |  |
| Anterior cingulate gyrus and sulcus RH |  |  |  |
|  | Group | 0.28 | 0.593 |
|  | Time | 1.93 | 0.164 |
|  | Group x Time | 0.05 | 0.818 |
| Long insular gyrus and central sulcus of the insula LH |  |  |  |
|  | Group | 0.73 | 0.391 |
|  | Time | 4.23 | 0.039 * |
|  | Group x Time | 0.02 | 0.882 |
| Medial orbital sulcus RH |  |  |  |
|  | Group | 0.99 | 0.317 |
|  | Time | 7.92 | 0.004 ** |
|  | Group x Time | 0.09 | 0.755 |
| Pallidum RH |  |  |  |
|  | Group | 2.05 | 0.151 |
|  | Time | 0.41 | 0.521 |
|  | Group x Time | 0.03 | 0.862 |
| Superior parietal sulcus RH |  |  |  |
|  | Group | 0.06 | 0.804 |
|  | Time | 0.07 | 0.779 |
|  | Group x Time | 0.62 | 0.428 |

*Note.* Results of the main effects and interactions from the mixed model testing for whether regions identified with predicting youth with rapid weight gain at the one-year time point (Adise et al., 2021)^3^ exhibit structural changes in response to sustained, two-year, weight gain onset. Effects are independent of BMI, age, sex, baseline puberty, race/ethnicity, and highest household education and random effects (i.e., scanner and subject change). Reference variables for categorical variables: Healthy Weight, Weight Stable (WS_HW_), Male, White, and Bachelor’s Degree. Time was not corrected for multiple comparisons because it was not an effect of interest but is reported for reader interpretation. G=gyrus; S=sulcus; RH=right hemisphere; LH=left hemisphere. ROI labels correspond to the Destrieux atlas labels. *=*p*<0.05; **=*p*<0.01; ***=*p*<0.001. *p* values are derived from the *F*-statistic. ^a^ =survived correction for multiple comparisons (n_tests_=36) for group and Group x Time interactions.

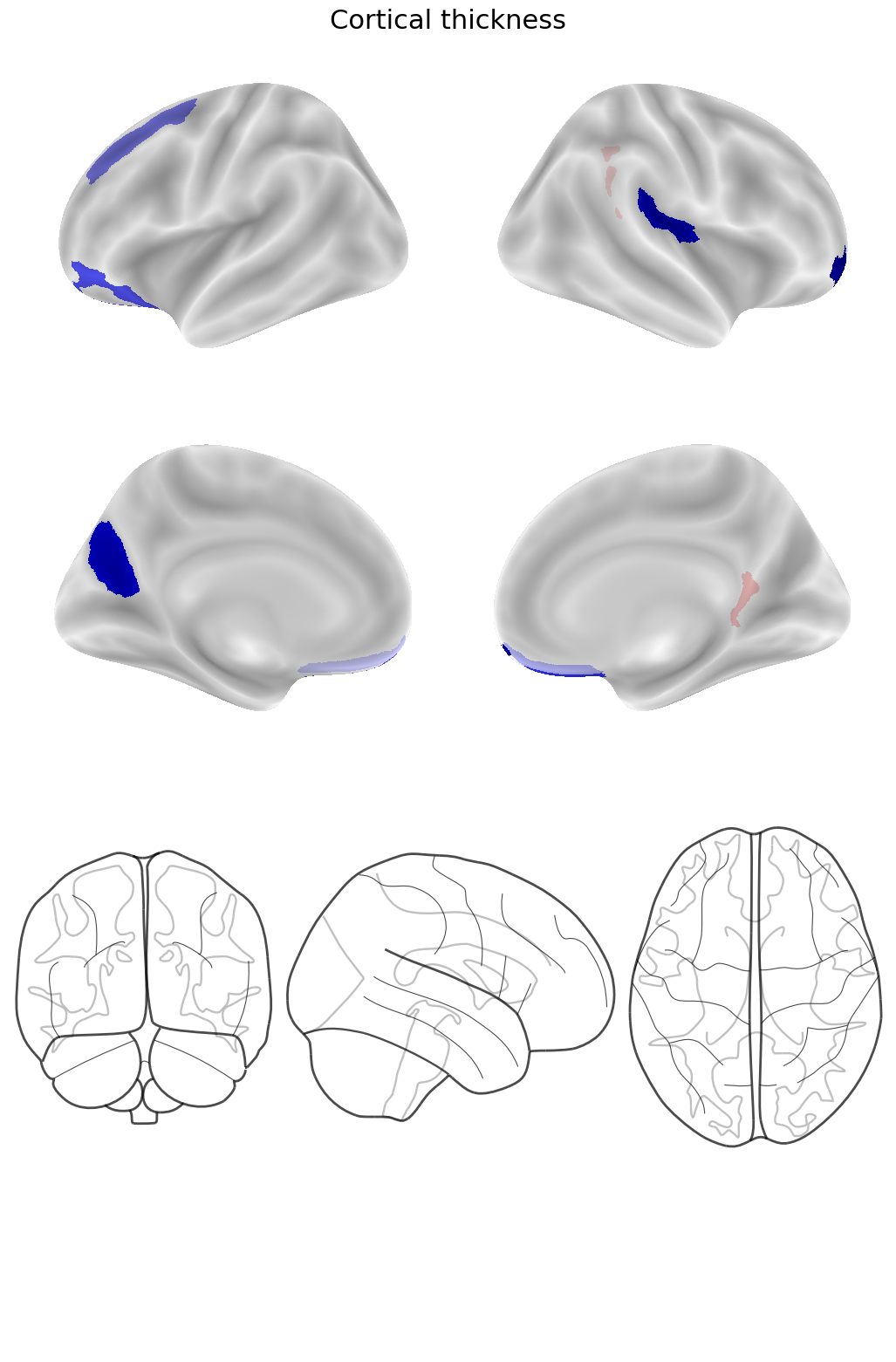

Cortical thickness

Surface area

Fractional anisotropy

Mean diffusivity

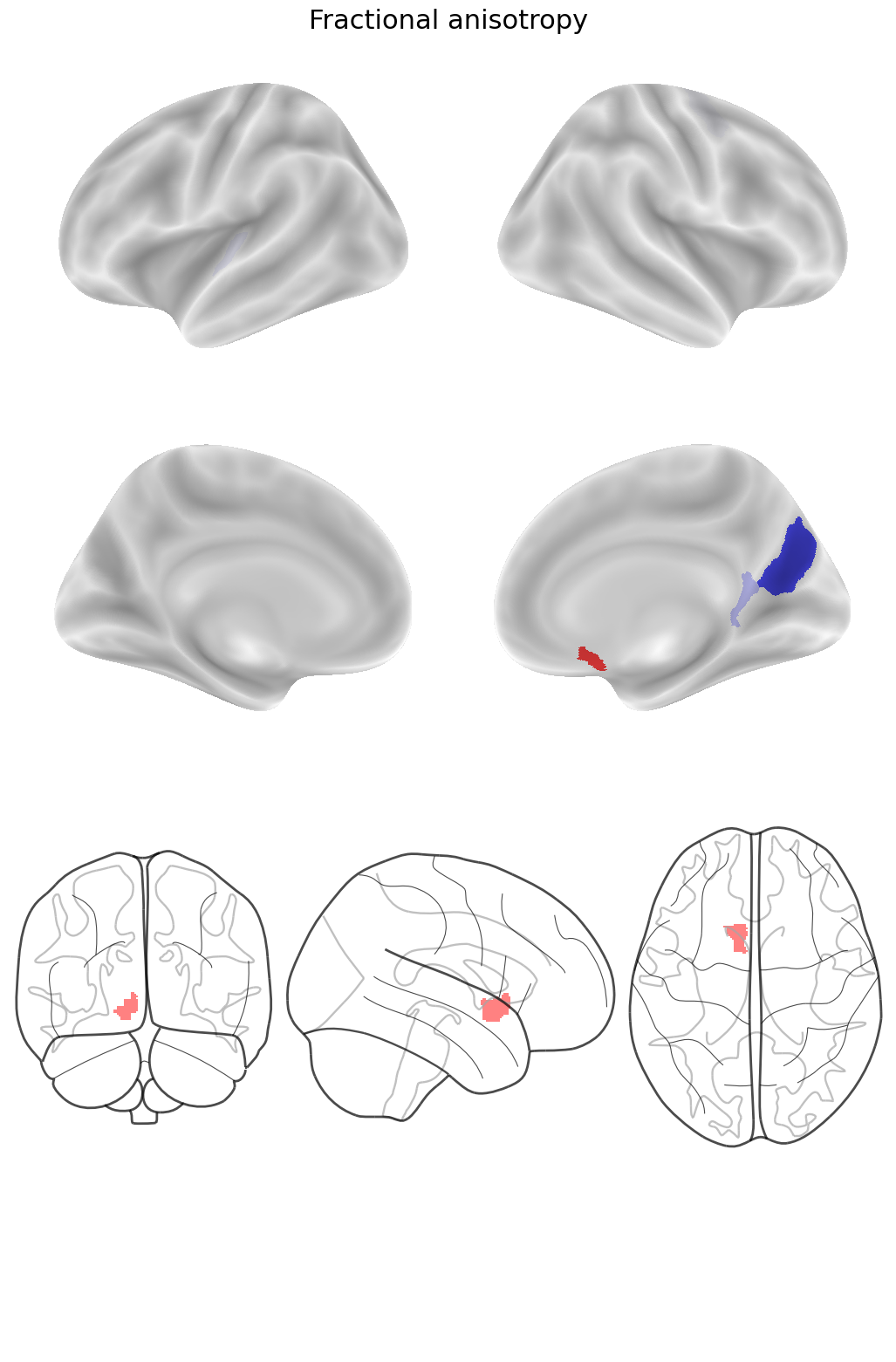

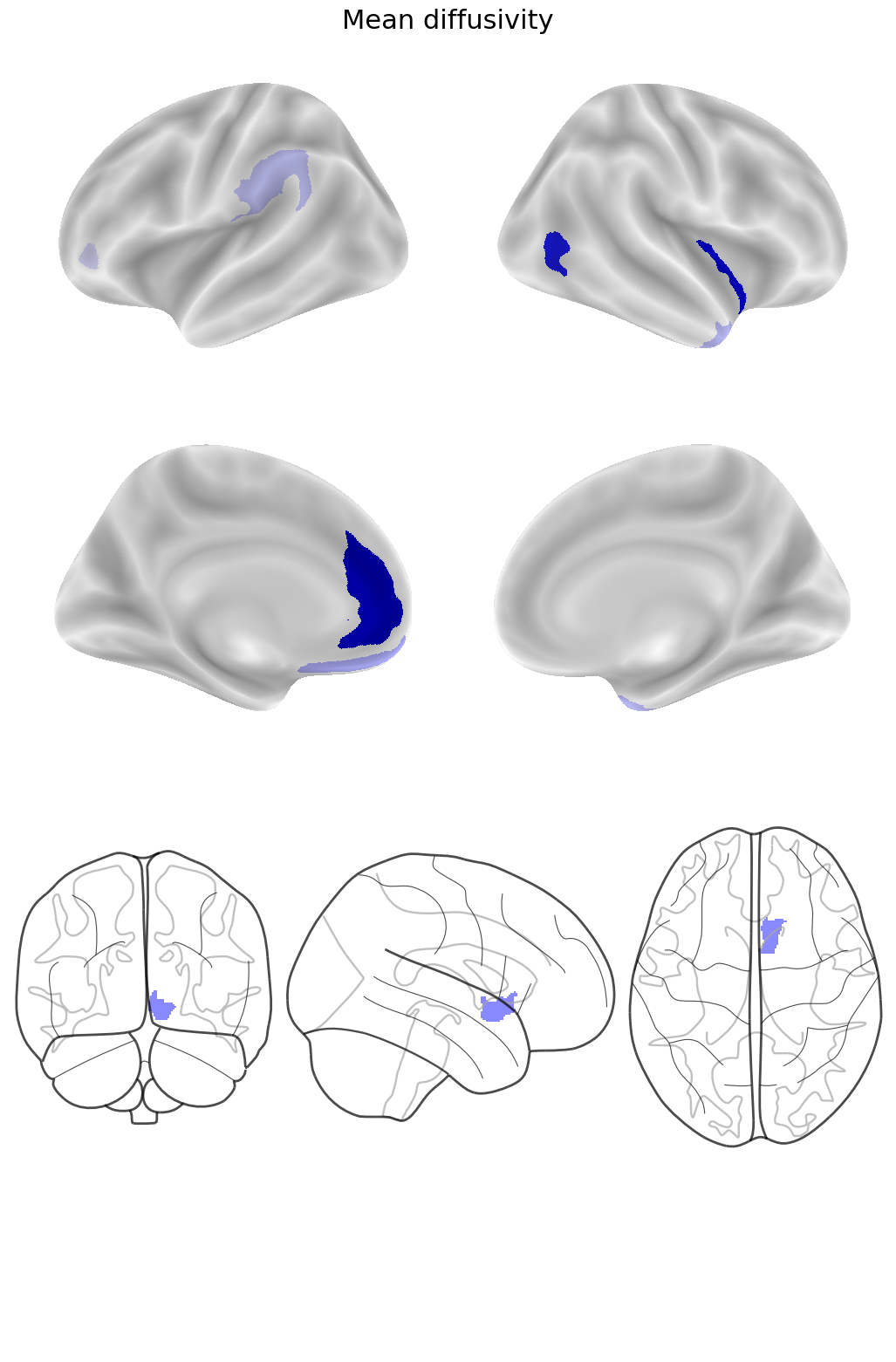

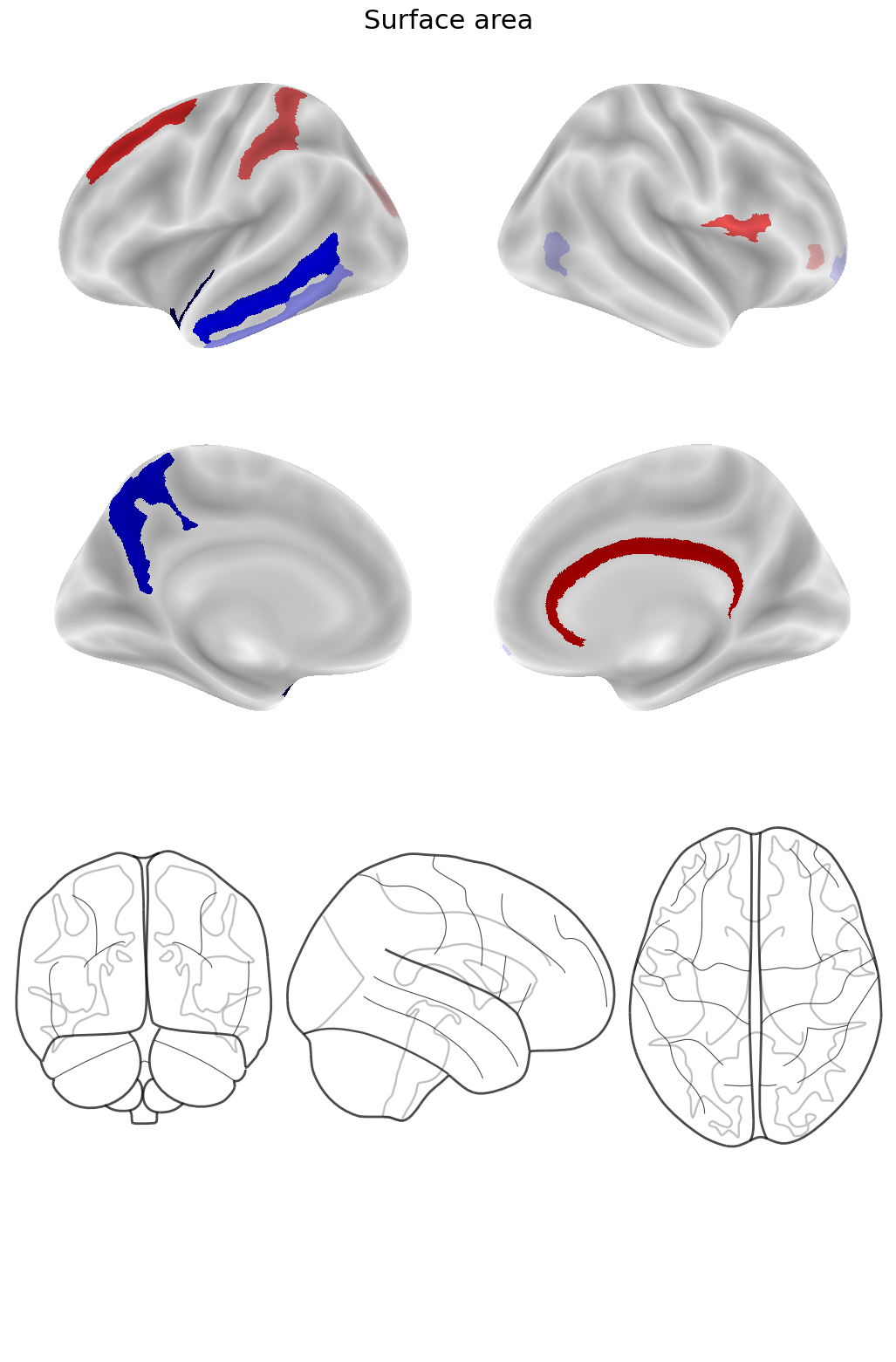

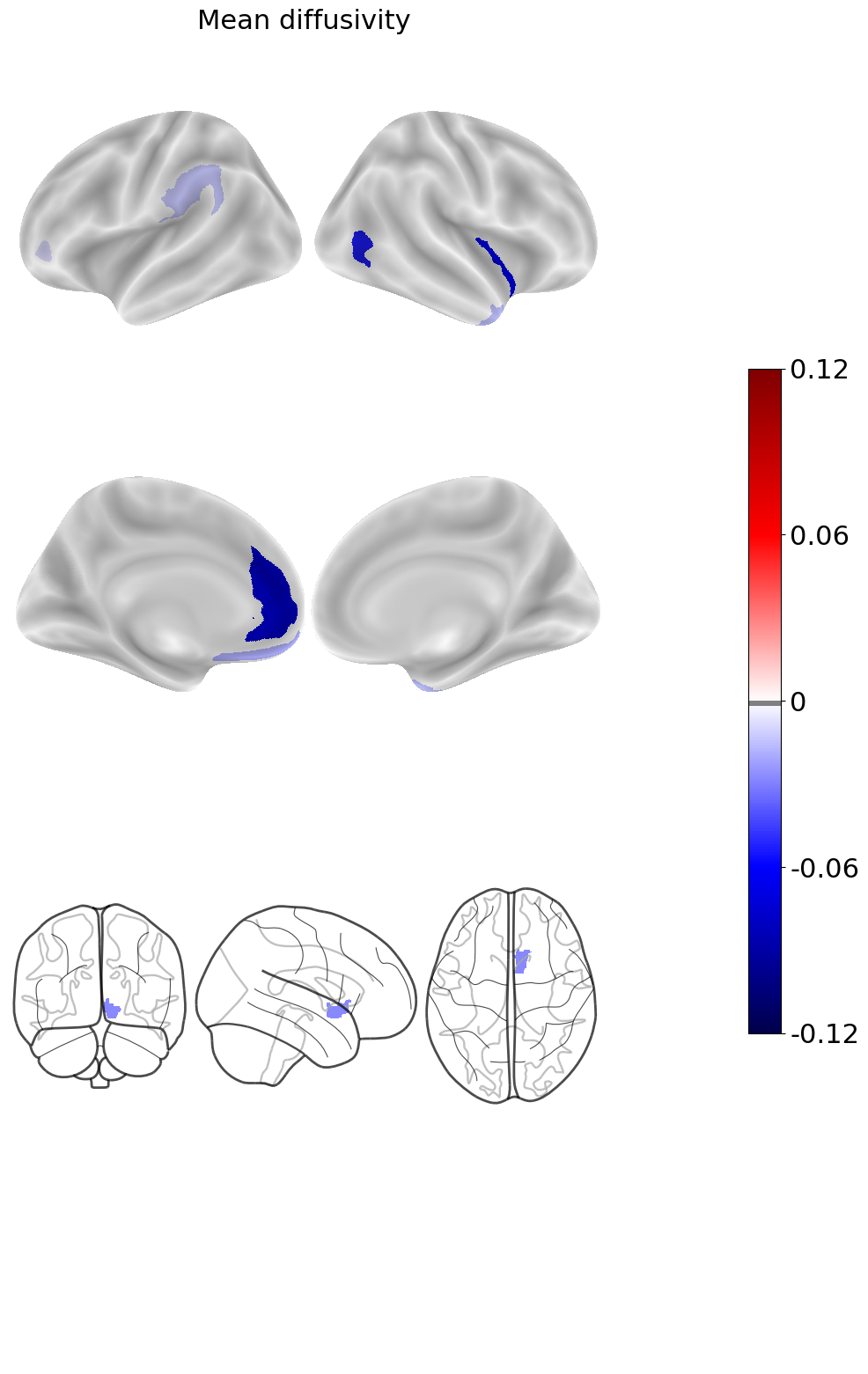

**Figure S2. V**isualization of the baseline brain features across each modality identified from the elastic net regression that predicted youth in the WG group after two-years of sustained, extreme weight gain. Colors correspond to the beta weights of the regression model.

**Table S6.** The model estimates for the training and test dataset for the prediction analysis in which baseline features were used to predict group membership at two-years.

|  | Training (n = 605, n _WG_ = 187) | | | Testing (n = 142, n _WG_ = 33) | | |  |
| --- | --- | --- | --- | --- | --- | --- | --- |
| Model | AUC | MCC | Bal Acc | AUC | MCC | Bal Acc | # of features |
| Brain only | 0.68 | 0.22 | 0.61 | 0.65 | 0.20 | 0.61 | 240 |
| Nonbrain only | 0.74 | 0.32 | 0.67 | 0.77 | 0.45 | 0.76 | 37 |
| Brain + Nonbrain | 0.76 | 0.38 | 0.70 | 0.72 | 0.23 | 0.63 | 44 |

*Note*. The area under the curve (AUC), Matthews Correlation Coefficient (MCC), and balanced accuracy (Bal acc) as well as the number of features selected from the model are presented. Covariates were dummy coded for Race, sex, highest household education, and MRI scanner serial number. The brain features included region of interest (ROI) estimates for cortical thickness, surface area, diffusion tensor imaging (DTI) estimates of fractional anisotropy (FA) and mean diffusivity (MD), and subcortical volume regions. WG = weight gain.

**Table S7.** Results of the main effects and interactions from the mixed model testing for whether regions identified at baseline as predictive of WG at the two-year period exhibited structural changes in response to WG onset.

| **ROI** |  | ***F*** | ***p*** |
| --- | --- | --- | --- |
| *Cortical Thickness* |  |  |  |
| Frontomarginal gyrus RH |  |  |  |
|  | Group | 19.2 | <0.001 ^a^ *** |
|  | Time | 2.18 | 0.140 |
|  | Group x Time | 9.86 | 0.001 ^a^ *** |
| Posterior ventral cingulate gyrus RH |  |  |  |
|  | Group | 3.08 | 0.079 . |
|  | Time | 1.17 | 0.277 |
|  | Group x Time | 0.00 | 0.930 |
| Orbital gyrus LH |  |  |  |
|  | Group | 3.85 | 0.049 * |
|  | Time | 2.04 | 0.152 |
|  | Group x Time | 0.48 | 0.485 |
| Rectal gyrus LH |  |  |  |
|  | Group | 1.87 | 0.171 |
|  | Time | 0.07 | 0.783 |
|  | Group x Time | 0.31 | 0.574 |
| Rectal gyrus RH |  |  |  |
|  | Group | 9.21 | 0.002 ^a^ ** |
|  | Time | 2.75 | 0.097 . |
|  | Group x Time | 2.28 | 0.130 |
| Posterior ramus of the lateral sulcus RH |  |  |  |
|  | Group | 5.76 | 0.016 * |
|  | Time | 0.01 | 0.898 |
|  | Group x Time | 1.04 | 0.307 |
| Superior frontal sulcus LH |  |  |  |
|  | Group | 1.54 | 0.214 |
|  | Time | 2.49 | 0.998 |
|  | Group x Time | 0.08 | 0.770 |
| Sulcus intermedius primus (of Jensen) RH |  |  |  |
|  | Group | 0.11 | 0.731 |
|  | Time | 0.29 | 0.590 |
|  | Group x Time | 0.24 | 0.620 |
| Medial orbito-olfactory sulcus RH |  |  |  |
|  | Group | 7.43 | 0.006 ** |
|  | Time | 0.17 | 0.676 |
|  | Group x Time | 0.46 | 0.495 |
| Parieto-occipital sulcus LH |  |  |  |
|  | Group | 6.05 | 0.014 * |
|  | Time | 1.47 | 0.224 |
|  | Group x Time | 0.10 | 0.743 |
| *Surface Area* |  |  |  |
| Frontomarginal gyrus RH |  |  |  |
|  | Group | 0.04 | 0.829 |
|  | Time | 0.04 | 0.825 |
|  | Group x Time | 0.09 | 0.757 |
| Inferior frontal opercular gyrus RH |  |  |  |
|  | Group | 0.34 | 0.555 |
|  | Time | 2.49 | 0.114 |
|  | Group x Time | 1.83 | 0.175 |
| Precuneus gyrus LH |  |  |  |
|  | Group | 0.49 | 0.481 |
|  | Time | 0.45 | 0.499 |
|  | Group x Time | 0.76 | 0.382 |
| Planum polare of the superior temporal gyrus LH |  |  |  |
|  | Group | 0.23 | 0.630 |
|  | Time | 0.24 | 0.617 |
|  | Group x Time | 2.86 | 0.090 . |
| Inferior temporal gyrus LH |  |  |  |
|  | Group | 5.05 | 0.024 * |
|  | Time | 2.44 | 0.118 |
|  | Group x Time | 0.87 | 0.349 |
| Middle temporal gyrus LH |  |  |  |
|  | Group | 0.22 | 0.636 |
|  | Time | 2.74 | 0.097 . |
|  | Group x Time | 1.33 | 0.247 |
| Inferior circula insula sulcus RH |  |  |  |
|  | Group | 0.29 | 0.586 |
|  | Time | 0.00 | 0.987 |
|  | Group x Time | 0.05 | 0.809 |
| Superior frontal sulcus LH |  |  |  |
|  | Group | 1.54 | 0.214 |
|  | Time | 0.37 | 0.538 |
|  | Group x Time | 0.18 | 0.663 |
| Superior occipital sulcus and transverse occipital sulcus LH |  |  |  |
|  | Group | 0.00 | 0.934 |
|  | Time | 0.02 | 0.868 |
|  | Group x Time | 4.42 | 0.035 * |
| Anterior occipital sulcus RH |  |  |  |
|  | Group | 0.15 | 0.698 |
|  | Time | 1.53 | 0.215 |
|  | Group x Time | 0.92 | 0.335 |
| Lateral orbital sulcus RH |  |  |  |
|  | Group | 2.74 | 0.097 . |
|  | Time | 2.11 | 0.145 |
|  | Group x Time | 2.64 | 0.104 |
| Pericallosal sulcus RH |  |  |  |
|  | Group | 1.12 | 0.288 |
|  | Time | 0.19 | 0.655 |
|  | Group x Time | 0.43 | 0.511 |
| Postcentral sulcus LH |  |  |  |
|  | Group | 2.38 | 0.123 |
|  | Time | 3.24 | 0.072 . |
|  | Group x Time | 0.00 | 0.968 |
| *Fractional Anisotropy* |  |  |  |
| Accumbens area LH |  |  |  |
|  | Group | 1.96 | 0.161 |
|  | Time | 43.4 | <0.001 *** |
|  | Group x Time | 1.75 | 0.186 |
| Posterior ventral cingulate gyrus RH |  |  |  |
|  | Group | 10.7 | 0.001 ^a^ ** |
|  | Time | 0.35 | 0.553 |
|  | Group x Time | 0.34 | 0.555 |
| Subcallosal gyrus RH |  |  |  |
|  | Group | 0.92 | 0.336 |
|  | Time | 17.1 | <0.001 *** |
|  | Group x Time | 2.45 | 0.117 |
| Anterior transverse temporal gyrus LH |  |  |  |
|  | Group | 3.42 | 0.064 . |
|  | Time | 0.36 | 0.546 |
|  | Group x Time | 0.66 | 0.415 |
| Parieto-occipital sulcus RH |  |  |  |
|  | Group | 10.2 | 0.001 ^a^ ** |
|  | Time | 0.42 | 0.514 |
|  | Group x Time | 0.11 | 0.736 |
| Superior part of the precentral sulcus RH |  |  |  |
|  | Group | 2.83 | 0.092 . |
|  | Time | 3.80 | 0.051 . |
|  | Group x Time | 1.41 | 0.235 |
| *Mean Diffusivity* |  |  |  |
| Accumbens area RH |  |  |  |
|  | Group | 15.4 | <0.001 ^a^ *** |
|  | Time | 22.6 | <0.001 *** |
|  | Group x Time | 2.07 | 0.150 |
| Anterior cingulate gyrus and sulcus LH |  |  |  |
|  | Group | 1.13 | 0.286 |
|  | Time | 0.39 | 0.530 |
|  | Group x Time | 0.00 | 0.923 |
| Long insular gyrus and central sulcus of the insula RH |  |  |  |
|  | Group | 1.45 | 0.227 |
|  | Time | 7.58 | 0.005 ** |
|  | Group x Time | 0.13 | 0.713 |
| Supramarginal gyrus LH |  |  |  |
|  | Group | 0.42 | 0.515 |
|  | Time | 0.14 | 0.706 |
|  | Group x Time | 0.01 | 0.890 |
| Rectal gyrus LH |  |  |  |
|  | Group | 0.05 | 0.817 |
|  | Time | 6.54 | 0.010 * |
|  | Group x Time | 1.25 | 0.263 |
| Subcallosal gyrus LH |  |  |  |
|  | Group | 4.18 | 0.041 * |
|  | Time | 15.8 | <0.001 *** |
|  | Group x Time | 0.21 | 0.642 |
| Temporal pole RH |  |  |  |
|  | Group | 4.67 | 0.030 * |
|  | Time | 0.02 | 0.883 |
|  | Group x Time | 0.33 | 0.560 |
| Superior circular insula sulcus LH |  |  |  |
|  | Group | 1.69 | 0.192 |
|  | Time | 0.24 | 0.618 |
|  | Group x Time | 0.49 | 0.482 |
| Anterior occipital sulcus RH |  |  |  |
|  | Group | 1.72 | 0.189 |
|  | Time | 0.22 | 0.633 |
|  | Group x Time | 2.80 | 0.094 . |
| Lateral orbital sulcus LH |  |  |  |
|  | Group | 0.43 | 0.507 |
|  | Time | 0.03 | 0.849 |
|  | Group x Time | 1.11 | 0.291 |

*Note.* Effects are independent of BMI, age, sex, baseline puberty, race/ethnicity, and highest parental education and random effects (e.g., scanner and subject). Reference variables for categorical variables: Healthy Weight, Weight Stable (WS_HW_), male, White, and Bachelor’s Degree. Time was not corrected for multiple comparisons because it was not an effect of interest but is reported for reader interpretation. CI = confidence interval; G = gyrus; S = sulcus; RH = right hemisphere; LH = left hemisphere. ROI labels correspond to the Destrieux atlas labels. * = *p*<0.05; ** = *p*<0.01; *** = *p*<0.001. *p* values are derived from the *F*-statistic. ^a^ = survived correction for multiple comparisons (n_tests_ = 78) for Group and Group x Time interactions.

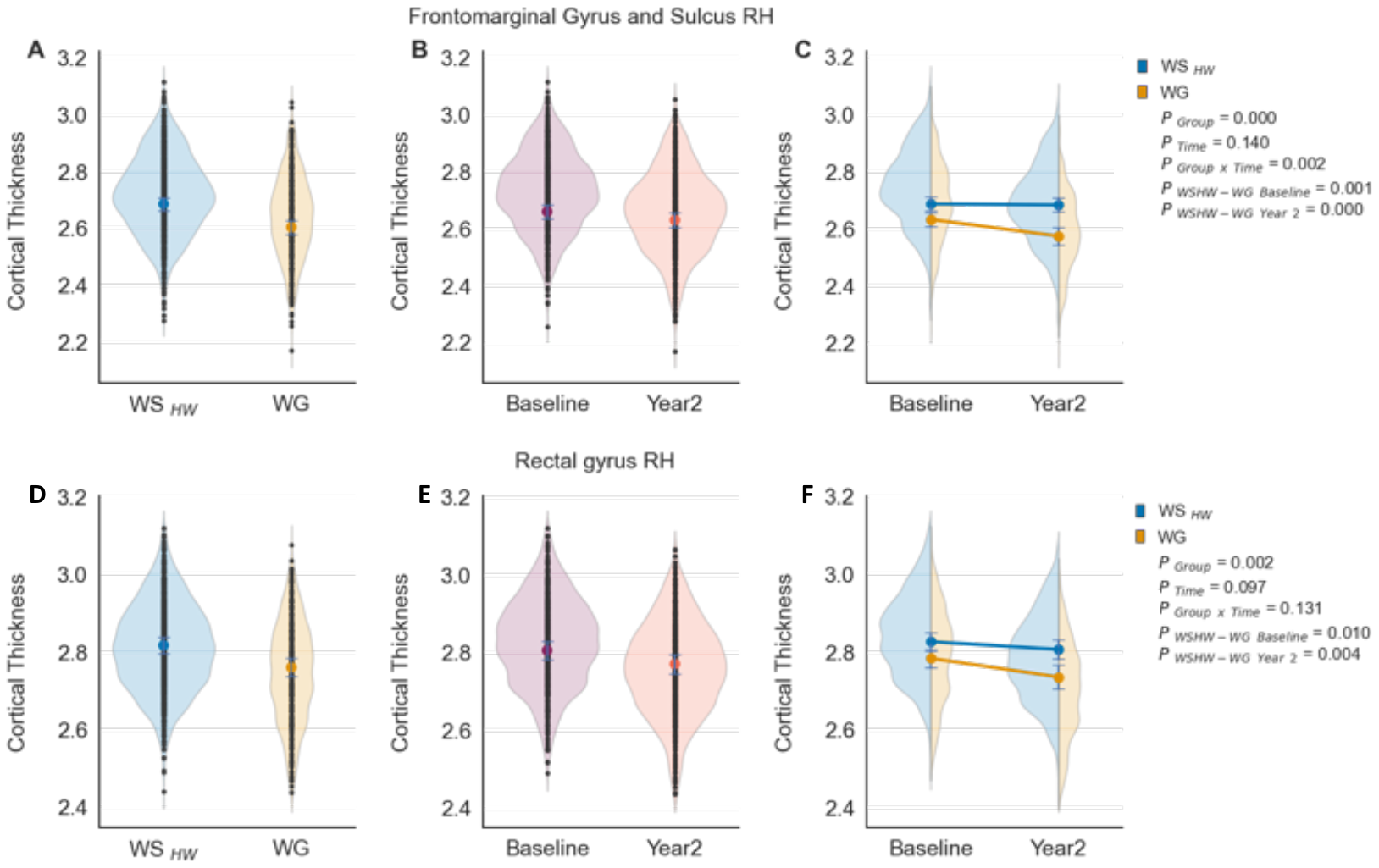
**Figure S3**. The distribution of estimates (i.e., violin plots and black dots) and means of the main effects and interactions adjusted for all covariates (e.g., age, sex, baseline puberty, BMI, race/ethnicity, highest household education, and caregiver report of prenatal exposure to alcohol and tobacco) and random effects (e.g., scanner and subject ID). **A** and **D)** Main effects of Group (Healthy Weight, Weight Stable [WS_HW_], Weight Gain [WG]) collapsed across time (baseline, year 2). **B** and **E)** Main effects of Time collapsed across group. **C** and **F)** Interactions between Group (WS_HW_ vs. WG) by Time (baseline vs. year 2). RH = right hemisphere

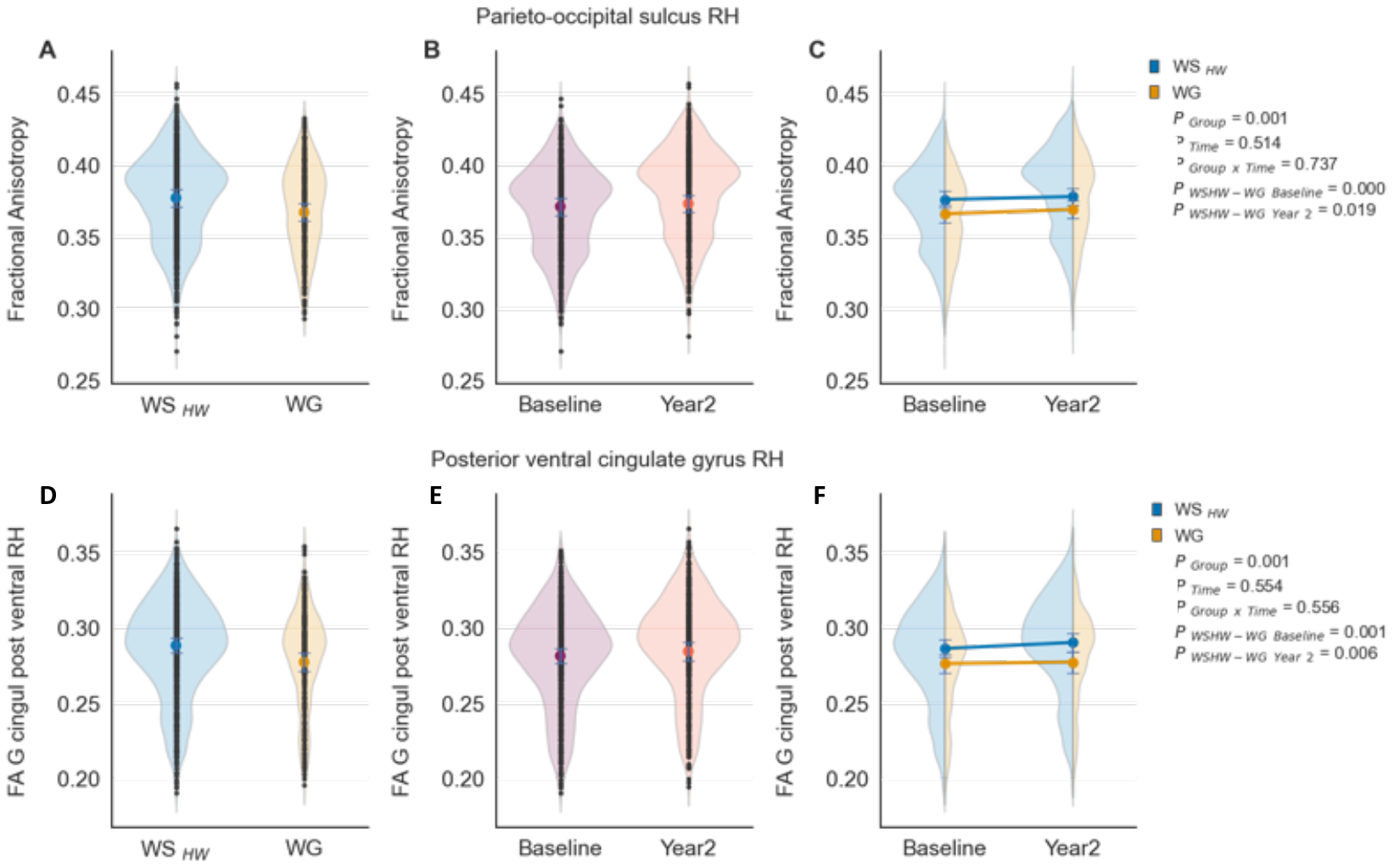

**Figure S4**. The distribution of estimates (i.e., violin plots and black dots) and means of the main effects and interactions adjusted for all covariates (e.g., age, sex, puberty, BMI, race/ethnicity, highest household education, and caregiver report of prenatal exposure to alcohol and tobacco) and random effects (e.g., scanner and subject ID). **A** and **D)** Main effects of Group (Healthy Weight, Weight Stable [WS_HW_], Weight Gain [WG]) collapsed across time (baseline, year 2). **B** and **E)** Main effects of Time collapsed across group. **C** and **F)** Interactions between Group (WS_HW_ vs. WG) by Time (baseline vs. year 2). RH = right hemisphere

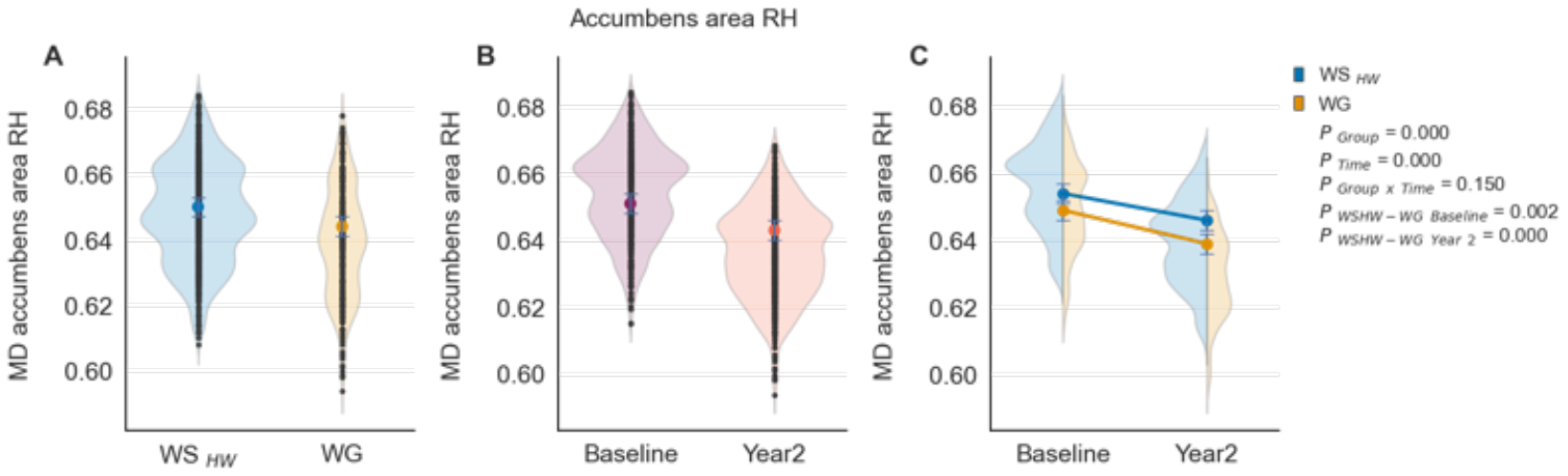

**Figure S5**. The distribution of estimates (i.e., violin plots and black dots) and means of the main effects and interactions adjusted for all covariates (e.g., age, sex, puberty, BMI, race/ethnicity, highest household education, and caregiver report of prenatal exposure to alcohol and tobacco) and random effects (e.g., scanner and subject ID). **A)** Main effects of Group (Weight Stable [WS_HW_], Weight Gain [WG]) collapsed across time (baseline, year 2). **B)** Main effects of Time collapsed across group. **C)** Interactions between Group (WS_HW_ vs. WG) by Time (baseline vs. year 2). RH = right hemisphere

**Table S8.** Post-hoc comparisons for the mixed effects models looking at group effects of brain structure change over time.

|  |  |  | **WG** | | | | **WS_HW_** | | |
| --- | --- | --- | --- | --- | --- | --- | --- | --- | --- |
| **Feature** | ***t*** | ***p*** | | ***M*** | | **95% CI** | ***M*** | **95% CI** | |
| *Cortical thickness* |  |  | |  | |  |  |  | |
| Frontomarginal gyrus RH | 4.4 | <0.001 | | 2.602±0.025 | | [2.545, 2.66] | 2.684±0.023 | [2.631, 2.736] | |
| Rectal gyrus RH | 3.04 | 0.001 | | 2.759±0.024 | | [2.704, 2.814] | 2.816±0.022 | [2.766, 2.866] | |
| *Fractional anisotropy* |  |  | |  | |  |  |  | |
| Parieto-occipital sulcus RH | 3.66 | <0.001 | | | 0.367±0.006 | [0.354, 0.38] | 0.377±0.006 | | [0.365 ,0.39] |
| Posterior ventral cingulate gyrus RH | 3.31 | 0.001 | | | 0.277±0.006 | [0.265, 0.29] | 0.288±0.005 | | [0.276, 0.3] |
| *Mean diffusivity* |  |  | |  | |  |  |  | |
| Accumbens area RH | 4.42 | <0.001 | | 0.643±0.003 | | [0.637,0.65] | 0.65±0.003 | [0.644,0.657] | |

***Note.*** Results of the corrected main effects from the mixed model testing for whether regions identified at baseline as predictive of weight gain group membership at the two-year follow-up exhibited structural changes in response to weight gain onset. Effects are independent of BMI, age, sex, baseline puberty, race/ethnicity, highest household education, random effects (e.g., scanner and subject), and caregiver report of prenatal exposure to alcohol and tobacco. Reference variables for categorical variables: weight stable, male, White, and Bachelor’s degree. CI=confidence interval; RH=right hemisphere; ROI labels correspond to the Destrieux atlas labels. *=*p*<0.05; **=*p*<0.01; ***=*p*<0.001. *p* values are derived from the *t*-statistic. ^a^ =survived correction for multiple comparisons (n_tests_=78) for Group and Group x Time interactions.

**Table S9.** Results for the linear random mixed effects models for each subscale of the BIS/BAS questionnaire.

| **Measure** |  | ***F*** | ***p*** |
| --- | --- | --- | --- |
| *BIS/BAS* |  |  |  |
| drive |  |  |  |
|  | Group | 0.24 | 0.617 |
|  | Time | 1.60 | 0.204 |
|  | Group x Time | 0.48 | 0.488 |
| reward |  |  |  |
|  | Group | 0.00 | 0.982 |
|  | Time | 1.94 | 0.163 |
|  | Group x Time | 0.08 | 0.771 |
| inhibition |  |  |  |
|  | Group | 2.08 | 0.148 |
|  | Time | 17.2 | < 0.001 *** |
|  | Group x Time | 3.14 | 0.076 . |
| fun seeking |  |  |  |
|  | Group | 0.65 | 0.416 |
|  | Time | 5.71 | 0.016 * |
|  | Group x Time | 1.06 | 0.303 |

*Note.* Group = Weight Stable (WS_HW_) versus Weight Gain (WG). Time = Baseline vs. Year 2. Covariates entered into the model were age, sex, BMI, baseline puberty, race/ethnicity, and parental highest education. Categorical variables consisted of sex, race/ethnicity, and parental highest education. Models were adjusted for random effects of scanner (i.e., site) and subject. ^a^ = survived correction for multiple comparisons (n_tests_ = 8) for group and Group x Time interactions.

**Table S10.** Results for the linear random mixed effects models for each subscale of the UPPS-P questionnaire.

| **Measure** |  | ***F*** | ***p*** |
| --- | --- | --- | --- |
| *UPPS-P* |  |  |  |
| Negative urgency |  |  |  |
|  | Group | 0.11 | 0.739 |
|  | Time | 17.2 | < 0.001 *** |
|  | Group x Time | 0.91 | 0.337 |
| Lack of planning |  |  |  |
|  | Group | 3.87 | 0.049 * |
|  | Time | 0.97 | 0.323 |
|  | Group x Time | 1.54 | 0.214 |
| Sensation seeking |  |  |  |
|  | Group | 0.21 | 0.646 |
|  | Time | 1.47 | 0.224 |
|  | Group x Time | 0.28 | 0.590 |
| Positive urgency |  |  |  |
|  | Group | 0.00 | 0.936 |
|  | Time | 11.5 | < 0.001 *** |
|  | Group x Time | 10.9 | < 0.001 ^a^ *** |
| Lack of perseverance |  |  |  |
|  | Group | 7.61 | 0.005 ^a^ ** |
|  | Time | 1.84 | 0.174 |
|  | Group x Time | 3.43 | 0.064 . |

Results for the linear random mixed effects models for each subscale of the UPPS-P questionnaire. Group=Weight Stable (WS_HW_) vs. Weight Gain (WG). Time=Baseline vs. Year 2. Covariates entered into the model were age, sex, BMI, baseline puberty, race/ethnicity, highest household education, and caregiver report of prenatal exposure to alcohol and tobacco. Categorical variables consisted of sex, race/ethnicity, highest household education, and caregiver report for prenatal alcohol and tobacco. Models were adjusted for random effects of scanner (i.e., site) and subject. ^a^ =survived correction for multiple comparisons (n_tests_=10) for group and Group x Time interactions.

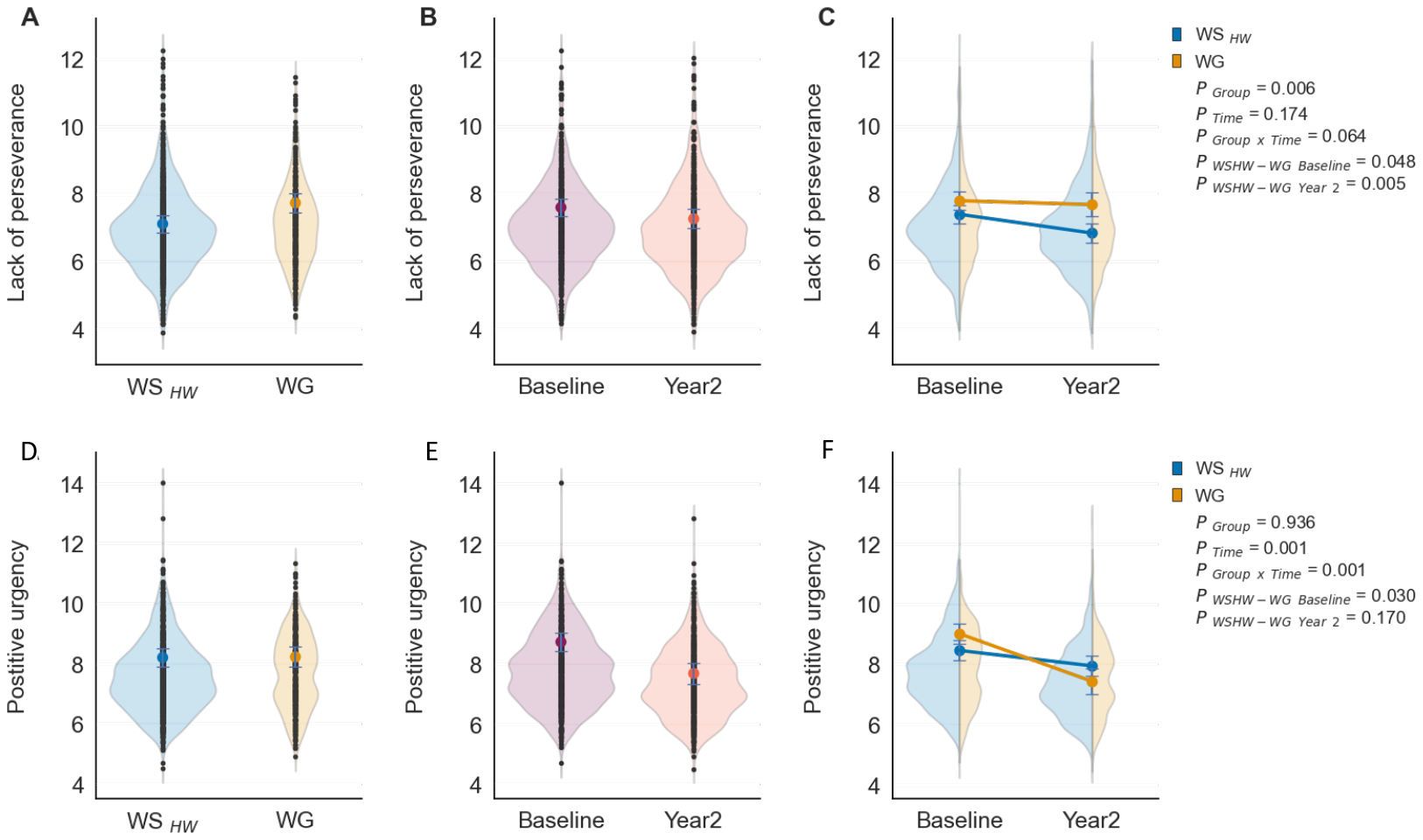
 **Figure S6.** The distribution of estimates (i.e., violin plots) and means (i.e., circles) for the positive urgency subscale of the UPPS-P. Positive urgency is the tendency to be impulsive during positive affect states. **A)** Main effect of Group (i.e., Weight Stable [WS_HW_] vs. Weight Gain [WG]) collapsed across time (i.e., baseline vs. year 2); **B)** Main effect of time collapsed acrossGgroup; **C)** Interaction between Group x Time. Effects were independent of age, sex, BMI, puberty, race/ethnicity, highest household education, and caregiver report of prenatal exposure to alcohol and tobacco . For **A** and **B**, the circle and error bars depict that marginal means and standard errors for the WS_HW_ and WG groups collapsed across time point (i.e., baseline to year 2). The black dots represent individual subject fit estimates. For **C**, the lines connecting the means indicate the change from baseline to year 2.
